# Activation of the antiviral factor RNase L triggers translation of non-coding mRNA sequences

**DOI:** 10.1101/2020.09.10.291690

**Authors:** Agnes Karasik, Grant D. Jones, Andrew V. DePass, Nicholas R. Guydosh

## Abstract

Ribonuclease L (RNase L) is activated as part of the innate immune response and plays an important role in the clearance of viral infections. When activated, it endonucleolytically cleaves both viral and host RNAs, leading to a global reduction in protein synthesis. However, it remains unknown how widespread RNA decay, and consequent changes in the translatome, promote the elimination of viruses. To study how this altered transcriptome is translated, we assayed the global distribution of ribosomes in RNase L activated human cells with ribosome profiling. We found that RNase L activation leads to a substantial increase in the fraction of translating ribosomes in ORFs internal to coding sequences (iORFs) and ORFs within 5’ and 3’ UTRs (uORFs and dORFs). Translation of these alternative ORFs was dependent on RNase L’s cleavage activity, suggesting that mRNA decay fragments are translated to produce short peptides that may be important for antiviral activity.

## INTRODUCTION

The detection of double stranded RNA (dsRNA) is a critical mechanism by which cells sense and defend against viral infections. In the cell, dsRNA activates several pathways important for type I interferon production, host translational shut down, and ultimately viral clearance or apoptosis (Hartmann, 2017; Kang et al., 2004; Malathi et al., 2007; Yoneyama et al., 2004). RNase <u>L</u> (Ribonuclease <u>L</u>atent), an endonuclease that broadly targets host and viral RNAs at UNN motifs (Han et al., 2014), is an important part of this antiviral defense network that is constitutively expressed in an inactive (monomer form in differentiated cells (Floyd-Smith et al., 1982; Zhou et al., 2005). Activation of RNase L during viral infections begins when <u>O</u>ligo<u>A</u>denylate <u>S</u>ynthetases (OASs) bind to and are activated by dsRNA. Once active, OASs synthesize 2-5-oligoadenylate (2-5A) from ATP (Li et al., 2016; Poulsen et al., 2015). Binding of 2-5A to RNase L stabilizes the dimeric form of the protein and, in turn, activates the enzyme by bringing the catalytic endoribonuclease domains into close proximity (Dong and Silverman, 1995; Han et al., 2014; Huang et al., 2014). Active RNase L is thought to be beneficial to the organism because it stimulates production of interferon (IFN), inhibits cell migration, and can trigger apoptosis of infected cells (Banerjee et al., 2015; Brennan-Laun et al., 2014; Castelli et al., 1997; Hassel et al., 1993; Zhou et al., 1997; Zhou et al., 1998).

Clearance of certain classes of viruses particularly depends on the activation of RNase L. For instance, mice lacking RNase L are not able to clear mouse encephalitis hepatitis virus, a member of the coronavirus family, resulting in higher mortality (Ireland et al., 2009). Furthermore, mutations in OAS genes are found in laboratory mice susceptible to West Nile Virus infection and in human patients with severe disease outcomes (Lucas et al., 2003; Mashimo et al., 2002; Yakub et al., 2005). While the activity of RNase L is thought to be antiviral, a role in bacterial infections is also emerging (Li et al., 2008). RNase L is also known to be a tumor suppressor in hereditary prostate cancer and other cancers (Carpten et al., 2002; Casey et al., 2002; Long et al., 2013; Wang et al., 2002) and was also identified as an oncogene in chronic myelogenous leukemia (Lee et al., 2013), further highlighting its role in human health.

Many aspects of how cleavage of host RNAs benefits the organism are unknown. RNase L cleaves a broad spectrum of host RNAs, including rRNAs, tRNAs, mRNAs and Y-RNAs (Burke et al., 2019; Donovan et al., 2017; Rath et al., 2015; Rath et al., 2019). While cleavage of rRNA was shown to not inhibit the activity of the ribosome (Rath et al., 2019), two recent studies established that, on average, mRNA levels in the cell can be reduced up to ~90% due to RNase L (Burke et al., 2019; Rath et al., 2019). This global effect was termed 2-5A Mediated Decay (2-5AMD), analogous to Nonsense Mediated Decay (NMD) (Rath et al., 2019).

Some antiviral mRNAs, such as IFN-β, were shown to be somewhat less sensitive to repression by RNase L activation. This was proposed to result from transcriptional compensation or lack of the most favorable cleavage sites (generally UA and UU motifs) (Burke et al., 2019; Floyd-Smith et al., 1981; Rath et al., 2015; Rath et al., 2019). It has also been reported that mRNAs encoding ribosomal proteins or miRNA binding sites are specifically targeted (Andersen et al., 2009; Rath et al., 2015). RNase L could therefore be important for generally reshaping the transcriptome and translatome to enhance the response to viral infection. Intriguingly, the RNase L cleavage fragments themselves have been proposed to be important since they can activate dsRNA sensors, such as RIG-I, MDA5, and PKR, that lead to IFN production or shutdown of translation initiation (Malathi et al., 2007; Manivannan et al., 2020). It remains unknown whether the fragments themselves have additional roles, such as the ability to be translated into short peptides.

There is some evidence that RNase L activation modulates translation. Early studies identified <u>R</u>Nase <u>L</u> <u>I</u>nhibitor (RLI) as a protein that binds RNase L. RLI was later shown to be the large subunit (60S) ribosome recycling factor, <u>A</u>TP <u>B</u>inding <u>C</u>assette sub-family <u>E</u> member 1 (ABCE1) (Bisbal et al., 1995; Khoshnevis et al., 2010; Pisarev et al., 2010; Young et al., 2015). Activated RNase L was also shown to bind to a translation termination factor, <u>e</u>ukaryotic <u>R</u>elease <u>F</u>actor 3 (eRF3) (Le Roy et al., 2005). Dual luciferase reporters in cell lysate suggest that activation of RNase L can induce translation of 3’ <u>U</u>n<u>T</u>ranslated <u>R</u>egions (UTRs) (Le Roy et al., 2005), an outcome that could be explained if RNase L inactivated these factors.

To answer the question of how RNase L activation impacts translation, we performed ribosome profiling on RNase L activated A549 human lung epithelial cells. We measured changes in ribosome distribution across the transcriptome and found RNase L activation leads to a shift in the distribution of translating ribosomes to 3’ and 5’ UTRs and internal <u>O</u>pen <u>R</u>eading <u>F</u>rames (ORFs) within the coding sequence. This alternative translation was dependent on the cleavage activity of RNase L and therefore suggests that RNA fragments can be translated, leading to the production of short peptides.

## RESULTS

### Host defense mRNAs are translated during RNase L activation

Since RNase L dramatically reshapes the transcriptome by degrading mRNA (Burke et al., 2019; Rath et al., 2019) and may itself affect the translation process, we investigated the distribution of ribosomes in cells where RNase L was active by performing ribosome profiling (Ribo-seq) (Ingolia et al., 2009) (Figure 1A). Active RNase L was shown to cleave both 18S and 28S rRNAs at precise locations (Cooper et al., 2014; Rath et al., 2019; Wreschner et al., 1981). Activation of RNase L is therefore traditionally assessed by using an rRNA cleavage assay (Wreschner et al., 1981). This activation can be specifically achieved by transfection with purified 2-5A (prepared as described in Methods) or the double stranded RNA mimic, poly I:C, that also activates additional dsRNA sensors, such as PKR, RIG-I and MDA5 (Chitrakar et al., 2019; de Haro et al., 1996; Martinand et al., 1998). We performed most experiments in the A549 lung carcinoma cell line since these cells had demonstrated robust RNase L activation in previous studies (Burke et al., 2019; Chitrakar et al., 2019; Rath et al., 2019). We found that 4.5 hours treatment of wild type (WT) A549 cells with 1-10 μM 2-5A was sufficient to generate readily detectable rRNA cleavage products by using electrophoretic analysis (BioAnalyzer) of purified total RNA (Figure 1B). The observed cleavage patterns were very similar when we treated with poly I:C (0.25 μg/ml, 4.5 hours), and no rRNA cleavage was detected in RNase L KO cells (Figure 1B), confirming cleavage specificity to RNase L. Next, we performed ribosome profiling by purifying ribosomes over a sucrose cushion and selecting mRNA fragments (25-34 nt) that correspond to ribosome footprints from 2-5A treated (or untreated) WT and RNase L KO A549 cells (McGlincy and Ingolia, 2017). After library preparation and deep-sequencing, ribosome protected footprints (referred to as footprints throughout the manuscript) were aligned to the human transcriptome and analyzed further (see Methods). Ribosome footprints are plotted by 5’ ends, rather than by 3’ ends or overall coverage, throughout the manuscript to enhance analysis of reading frame.

**Figure 1.**
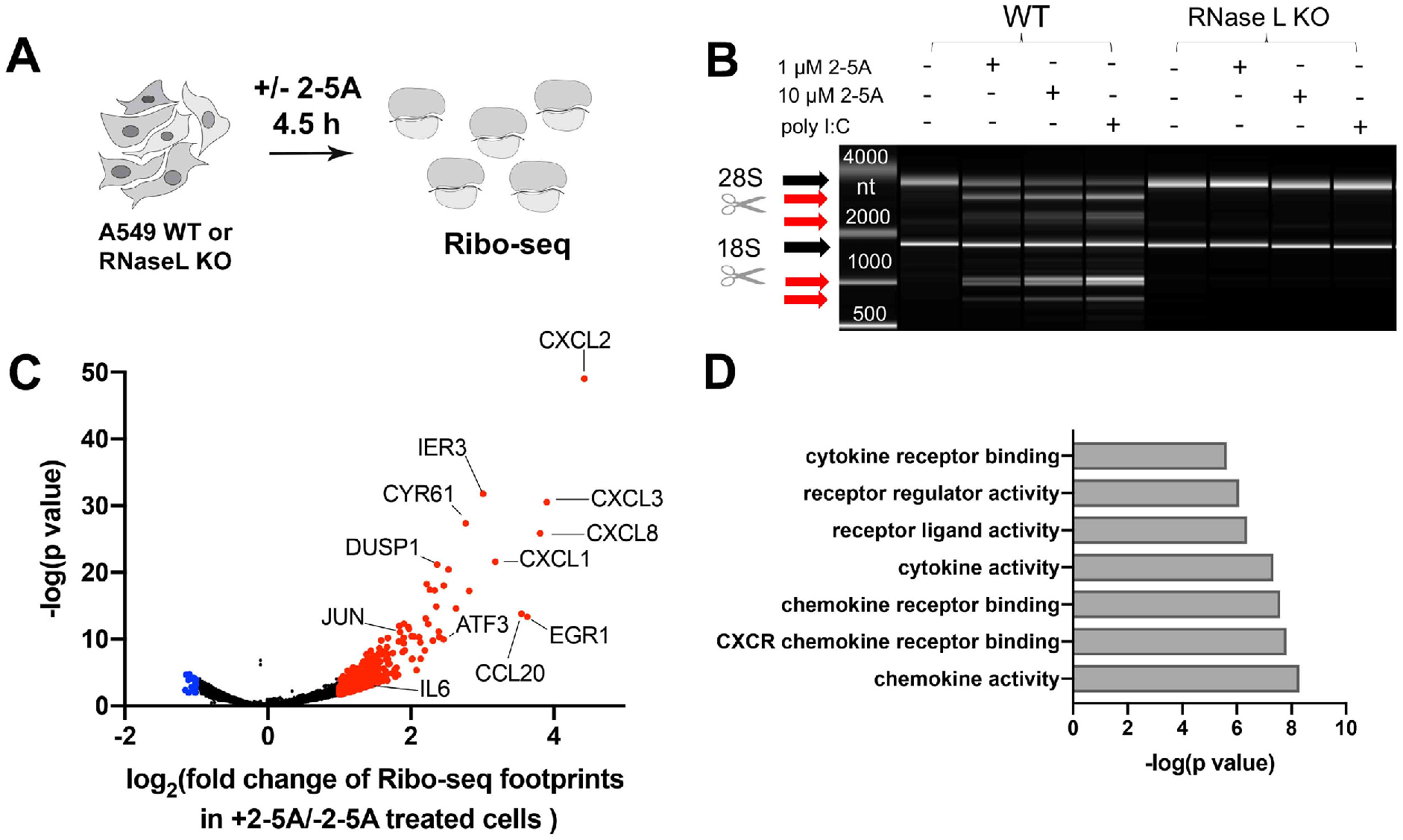
Changes in translation mirror known changes in mRNA levels. **A** Schematic representation of Ribo-seq experiments on 2-5 A treated or untreated A549 cells. First, cells were grown to ~70% confluency then 2-5 A was delivered by transfection. After 4.5 h, cells were harvested, and Ribo-seq experiments were performed as described in Methods. **B** BioAnalyzer-based rRNA cleavage assay showing activation of RNase L by 2-5 A and poly I:C in WT but not in RNase L KO A549 cells. Black arrows point to 28S and 18 S rRNA and red arrows indicate cleavage products. **C** Analysis of changes in ribosome footprint levels across coding sequences is consistent with previously-reported relative increases in immune-defense transcripts under RNase L activation. Data from two independent experiments were analyzed using DTEG.R (see more in Method Details). Genes with increased or decreased relative translation changes are marked red or blue, respectively (> or < 2-fold change in ribosome footprints and >0.05 p value). Genes related to immune response or analyzed later as examples are labeled. **D** Gene Ontology (GO) function analysis of transcripts with increased ribosome footprints in RNase L activated cells (>4-fold change, data in Table S1).

First, we investigated changes in translation during RNase L activation by analyzing the differences between ribosome footprint distribution in 2-5A treated and untreated cells. The mRNA pool is known to be greatly reduced by activation of RNase L (Burke et al., 2019; Rath et al., 2019). However, particular classes of mRNAs, including those involved in host defense against viruses, were shown to be somewhat resistant to RNase L-mediated degradation. Here, we employed ribosome profiling to evaluate whether ribosome occupancy levels also follow this trend when RNase L is active. We found significant changes in footprint distribution for many genes (344 genes >2-fold changes), including several pro-inflammatory chemokines and cytokines and regulators of immune response and cell physiology (Figure 1C and D). In particular, we found transcripts that were previously shown to increase in relative abundance during activation of RNase L show a similar relative increase in our ribosome profiling data, such as IL6 (interleukine-6) and EGR1 (Early growth response protein 1) (Burke et al., 2019). Thus, we confirm that differences in the abundance of ribosome footprints between RNase L activated and normal cells reflect changes in the abundance of these transcripts. We also noted that footprints from the genes encoding interferon-β and γ were not observed in 2-5A treated cells during the time course of the treatment, in agreement with previous findings (Burke et al., 2019). This absence suggests that pathways responsible for interferon production, such as those mediated by MDA-I and RIG-I, are not activated.

### Ribosomes accumulate in 3’UTRs during activation of RNase L

Since it was reported that RNase L activation increased translation of 3’UTR regions downstream of stop codons by interfering with factors that promote translation termination (eRF3) or ribosome recycling (ABCE1) (Le Roy et al., 2005), we assessed the level of ribosome footprints in 3’UTRs. This level was assayed by computing the ratio of footprints in every 3’UTR relative to its respective main ORF within the coding sequence (density ratio, 3’UTR:ORF) for each transcript (Young et al., 2015). We found that the ratios globally increased when WT A549 cells were treated with 2-5A, (Figure 2A, red dots above diagonal). This finding was consistent across several replicates as evidenced by boxplot analysis (Supplemental Figure 1A-C). We did not observe this trend in RNase L KO cells, showing that RNase L activation is required for the process. These data suggest that activation of RNase L increases translation of 3’UTRs relative to coding sequences.

**Figure 2.**
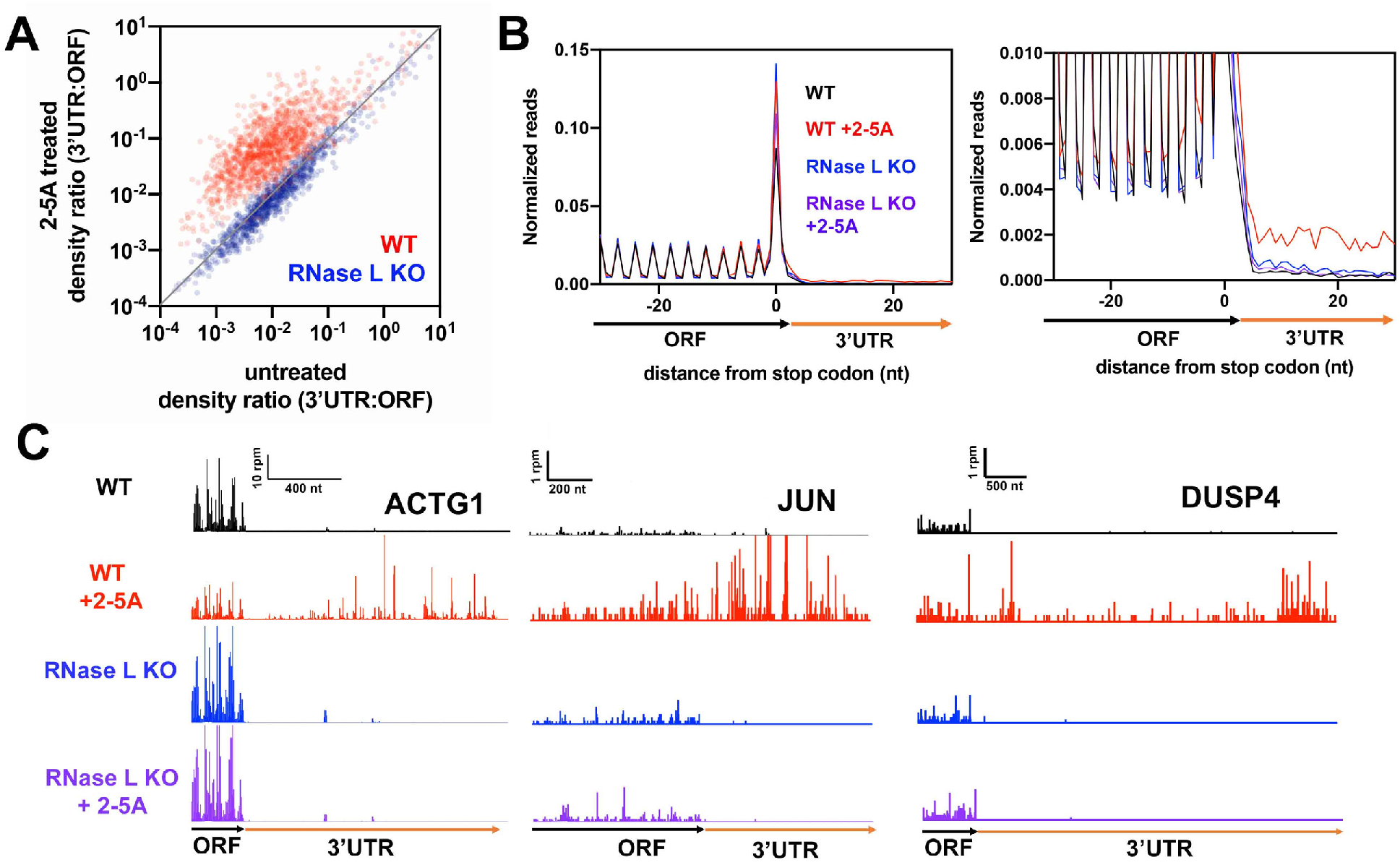
Activation of RNase L leads to increased ribosome footprint levels in 3’UTRs relative to coding regions. Related to Supplemental Figure 1. **A** Comparison of 3’UTR:ORF ribosome footprint density ratios in 2-5 A treated vs untreated conditions is shown for WT and RNase L KO A549 cells. Each dot represents one gene model. Genes plotted above the diagonal have increased relative 3’UTR translation in the treated compared to the untreated sample. **B** Normalized average ribosome footprint occupancy (metagene plot) around the stop codon of main ORFs reveals increased ribosome footprint levels in 3’UTRs. **C** Example gene models showing increased 3’UTR footprints in 2-5 A treated WT cells relative to untreated. Data from RNase L KO cells also shown as a control. Note that y-axis is truncated causing highest peaks in 3’UTRs to be higher than shown for JUN and ACTG1.

The increased presence of ribosomes in the 3’UTR was reproducible in other cell lines, such as Hap1 and HeLa, indicating that these observations are not cell line specific (Supplemental Figure 1D-G). Additionally, we tested whether the form of 2-5A used to activate RNase L modulated this observation. While the trimeric form of 2-5A we used for most experiments here is thought to be the shortest 2-5A molecule capable of activating RNase L, longer forms also achieve activation (Han et al., 2014; Le Roy et al., 2005). To verify that other forms of 2-5A don’t change the observed effect on translation in 3’UTRs, we tested whether non-trimeric forms of 2-5A (dimer and tetramer) have the same effect. We found that the tetramer also increased the 3’UTR:ORF density ratio, much as observed for the trimer form of 2-5A (in HeLa cells, Supplemental Figure 1E-G). In contrast, treatment with dimeric 2-5A did not induce rRNA cleavage, as expected (data not shown). It also did not increase relative translation of 3’UTRs, offering further evidence that the observed effects are dependent on the activation of RNase L (Supplemental Figure 1H and I).

To further investigate the mechanism by which RNase L activation was promoting translation of 3’UTRs, we aligned genes by their stop codons and averaged their respective ribosome footprints (“metagene” analysis). As expected, the average of footprints mapping to coding sequences (upstream of stop codons) showed strong three nucleotide (3-nt) periodicity, reflecting elongation by the ribosome across the main ORF (Figure 2B, left). In agreement with the 3’UTR:ORF density ratio analysis, we observed an increase in the average level of footprints in the 3’UTR for WT, but not for RNase L KO cells that were transfected with 2-5A (Figure 2B, right). In addition to this global analysis, individual analysis of 3’UTR regions from some transcripts, such as *JUN, ACTG* and *DUSP4*, revealed heavy translation upon RNase L activation (Figure 2C).

We then examined several properties of the 3’UTR average footprints to assess whether existing models (data reproduced in Supplemental Figure 1J) could explain how ribosomes bypass stop codons. It is feasible that lack of the termination factor (eRF3) would result in the ribosome “reading through” the stop codon and producing a C-terminally extended protein product by allowing the binding of a near-cognate tRNA and incorporation of an amino acid by the catalytic center at the stop codon. One of the hallmarks of “readthrough” of stop codons is that ribosome footprints in the 3’UTR maintain the reading frame established by the main ORF (Wangen and Green, 2020; Young et al., 2015; Young et al., 2018) (Supplemental Figure 1J). If activated RNase L triggered readthrough by inhibiting eRF3 via its proposed interaction (Le Roy et al., 2005), we would expect to observe 3-nt periodicity in the 3’UTR. However, the footprints that mapped to the 3’UTR region, on average, lacked this periodicity (Figure 2B, right). This suggests that the observed effects of RNase L activation are not caused by readthrough. Alternatively, loss of ribosome recycling activity due to lower levels of ABCE1 can increase average 3’UTR footprint levels in all 3 reading frames due to reinitiation of translation (Young et al., 2015). When ribosome recycling is inhibited, the encoded protein is released at the stop codon. However, because the unrecycled ribosome remains on the mRNA, it can move into the 3’UTR, translate downstream sequences, and produce short peptides. Since RNase L is known to directly interact with ABCE1 (Bisbal et al., 1995), it is possible that this interaction interferes with ABCE1’s 60S ribosome recycling function. However, the signature of lowered ABCE1 levels is an increased peak at the stop codon, due to the accumulation of unrecycled 80S ribosomes (Mills et al., 2016; Young et al., 2015) (Supplemental Figure 1J). However, stop codon peaks in the metagene analysis did not change in a way that supports this hypothesis, indicating that 2-5A treatment did not affect ribosome splitting (Figure 2B). In addition, other forms of recycling defect, such as loss of 40S subunit recycling, could explain the increased translation in 3’UTRs. However, defects in 40S recycling are known to result in ribosome queuing upstream of the stop codon (Young et al., 2018) (Supplemental Figure 1J), and this defect was not observed in RNase L activated cells (Figure 2B). More broadly, any translation in the 3’UTR that results from loss of termination or recycling activity should be characterized by a loss in footprint levels at distal regions of 3’UTRs since ribosomes will eventually terminate translation and be recycled in the 3’UTR, even if inefficiently (Young et al., 2015; Young et al., 2018) (Supplemental Figure 1J). However, a metagene plot computed from genes with long 3’UTRs revealed almost no decreasing trend on average and instead showed steady footprint levels over 1 kb of 3’UTR (Supplemental Figure 1K, compare to 1J). Metagene analysis therefore revealed that the increased translation in 3’UTRs did not mimic any known ribosome termination or recycling defect.

### RNase L activation induces translation of downstream ORFs (dORFs)

Since we observed increased relative ribosome occupancy in 3’UTRs, we next asked whether the ribosomes in 3’UTR regions were actively translating or nonproductively bound. We therefore looked for the hallmarks of active translation, such as peaks on start codons (indication of translation initiation) and 3-nt periodicity downstream from the dORF start codon (indication of elongation by ribosomes) in the 3’UTR. Consistent with a model where ribosomes in the 3’UTR were translating, we observed ribosome footprint peaks in 3’UTRs at canonical (AUG) or non-canonical (CUG, UUG and GUG) start codons in all three frames with respect to the main ORF. As an example, we show putative dORFs with the most prominent peaks in the 3’UTR of the ACTG1 mRNA (Figure 3A).To further confirm the existence of active translation, we extended the analysis globally by aligning 3’UTR start codons together and averaging ribosome footprints (metagene plots at dORFs). The dORF metagene analysis confirmed a large ribosome average footprint peak at canonical and non-canonical start codons. Importantly, this analysis also revealed 3-nt periodicity after AUG codons alone, consistent with AUG codons efficiently initiating translation that leads to elongation across downstream sequences (Figure 3B, Supplemental Figure 2). While we also observed a strong peak when non-canonical CUG start codons were separately analyzed, the 3-nt periodicity downstream was weaker than that in the dORF metagene analysis for AUG start codons (Figure 3C, Supplemental Figure 2). These trends are consistent with CUG being less efficient at initiating translation (Mehdi et al., 1990). In contrast, a similar analysis of untreated WT and treated or untreated RNase L KO cells did not show these characteristics, as expected (Figure 3B, Supplemental Figure 2).

**Figure 3.**
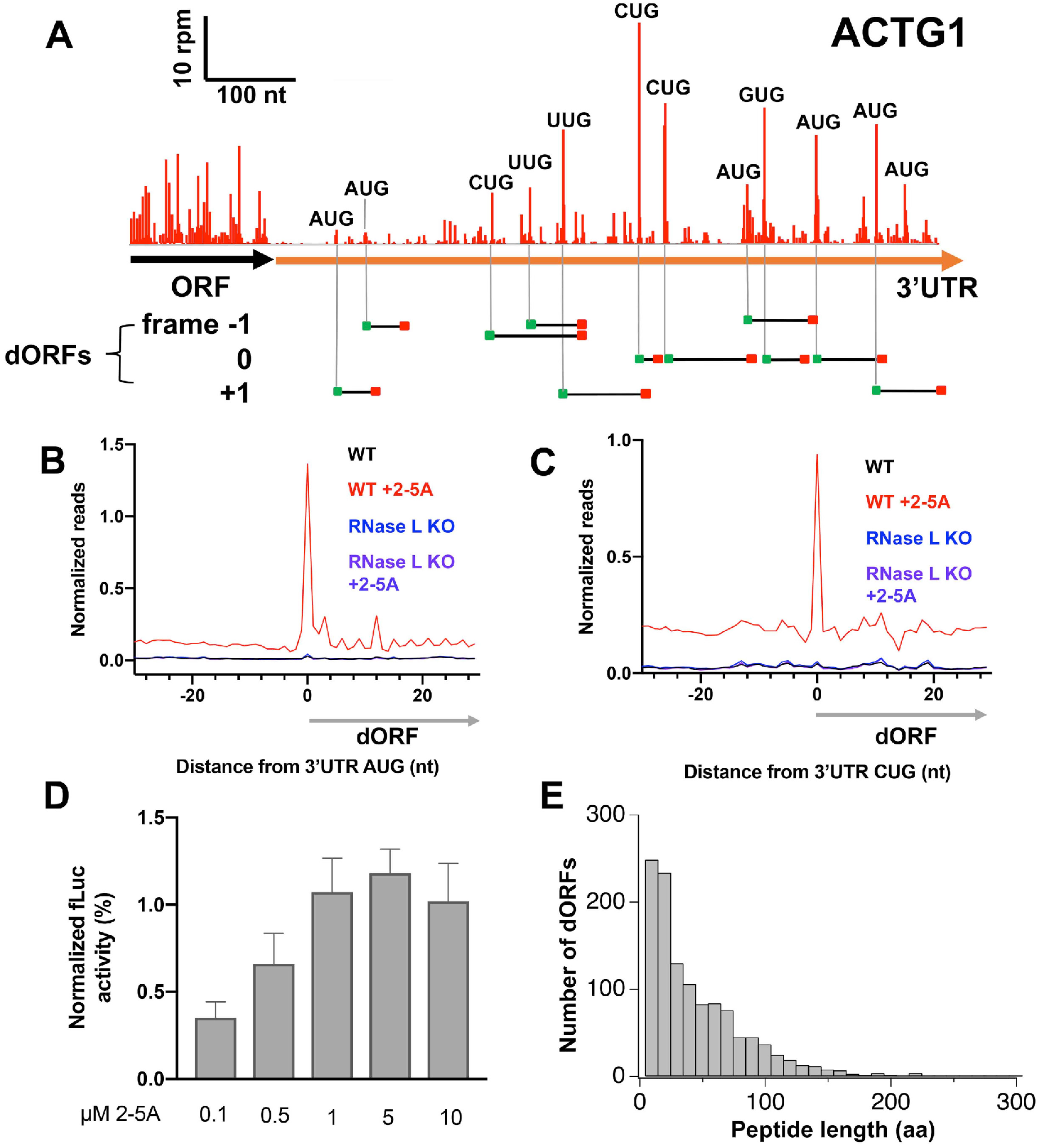
dORFs are translated during RNase L mediated antiviral response. See also Supplemental Figure 2. **A** Footprints that map to the 3’UTR of the ACTG1 gene model during RNase L activation. Cognate (AUG) and near-cognate (CUG, UUG and GUG) start codons are depicted as green squares and potential dORFs (black lines) are marked for prominent peaks that occur in all frames, signifying likely dORF translation events. Red squares represent the first stop codon after the start codon. **B** Average footprints around AUG start codons in 3’UTRs show an increased peak at the start codon followed by footprints that exhibit 3-nt periodicity in 2-5 A treated WT cells, but not in RNase L KO cells. Ribosome footprints are normalized to main ORF footprint levels prior to averaging. **C** Average ribosome footprint levels as in B, but for CUG start codons, also show an increased peak. **D** Results from *in vitro* dual luciferase assay where rLuc serves as main ORF and fLuc serves as a downstream ORF reveal the 2-5A concentration dependence of dORF translation. **E** Analysis of the length of dORF sequences that show increased ribosome footprint levels during RNase L activation. Analysis revealed 1192 peptides (>4 amino acid long and dORF with at least 5 aligned raw reads) as likely to be translated. Data were reproducible in all replicates. Data from analysis of replicate 1 are shown.

Despite these clear hallmarks of active dORF translation in RNase L activated cells, we wanted to further confirm that translation of these regions was occurring. We therefore employed a dual luciferase assay to directly measure the synthesis of a peptide encoded in the 3’UTR (Houck-Loomis et al., 2011). In this assay, the main ORF of the reporter gene encoded renilla luciferase (rLuc) and the coding sequence for firefly luciferase (fLuc) was inserted in the 3’UTR to act as a dORF (see Methods). Translation of the reporter was carried out in HeLa cell extracts in the presence or absence of different concentrations of 2-5A. Main ORF and dORF translation could therefore be measured by assaying for the activity of each respective luciferase enzyme. We found dORF fLuc activity increased relative to that of the main-ORF rLuc in a 2-5A concentration dependent manner (0.1-10 μM) (Figure 3D). Notably, this concentration dependence of 3’UTR translation was similar to that observed for the dimerization of an RNase L based 2-5A sensor, consistent with the idea that dORF translation is correlated with RNase L dimerization and activation (Chitrakar et al., 2019). This trend therefore provides added evidence that activation of RNase L increases translation of 3’UTR dORFs relative to the corresponding main ORF.

Taken together, ribosome profiling and a luciferase reporter assay showed that ribosomes translate dORFs when RNase L is activated. To offer further context about the possible magnitude of this effect, we computed the ribosome density across individual dORFs (initiated by AUG start codons, see Methods). Using conservative metrics for dORF translation (see legend), we found evidence for over a thousand dORFs being translated in response to RNase L activation. While most dORFs encode short peptides (less than 50 aa), many were found to encode longer products (Figure 3E).

### RNase L activation increases the translation of upstream ORFs (uORFs)

Since we observed increased dORF translation in RNase L activated cells without apparent defects in translation termination or recycling, we asked whether these seemingly anomalous translation events were also occurring in upstream ORFs (uORFs) in the 5’UTR. Most uORFs encode short peptides and start with an AUG or a near-cognate start codon. They are believed to act as regulators of downstream translation of coding sequences (Hinnebusch, 1993; Johnstone et al., 2016; Spealman et al., 2018) but may have other roles, such as regulating NMD or producing functional peptides (Chen et al., 2020; Hinnebusch, 1993; Johnstone et al., 2016; Lin et al., 2019). Therefore, we computed the ratio of ribosome footprint density between 5’UTRs and their respective main ORFs, similar to the 3’UTR analysis above. While basal translation of 5’UTRs in untreated cells is much higher than of 3’UTRs, we found that 5’UTR:ORF ratios were further increased in RNase L activated cells (Figure 4A, dots above diagonal, Supplemental 3A-C). As with 3’UTRs, the trend was eliminated in 2-5A treated cells lacking RNase L, indicating that the effect is attributable to RNase L activation.

**Figure 4.**
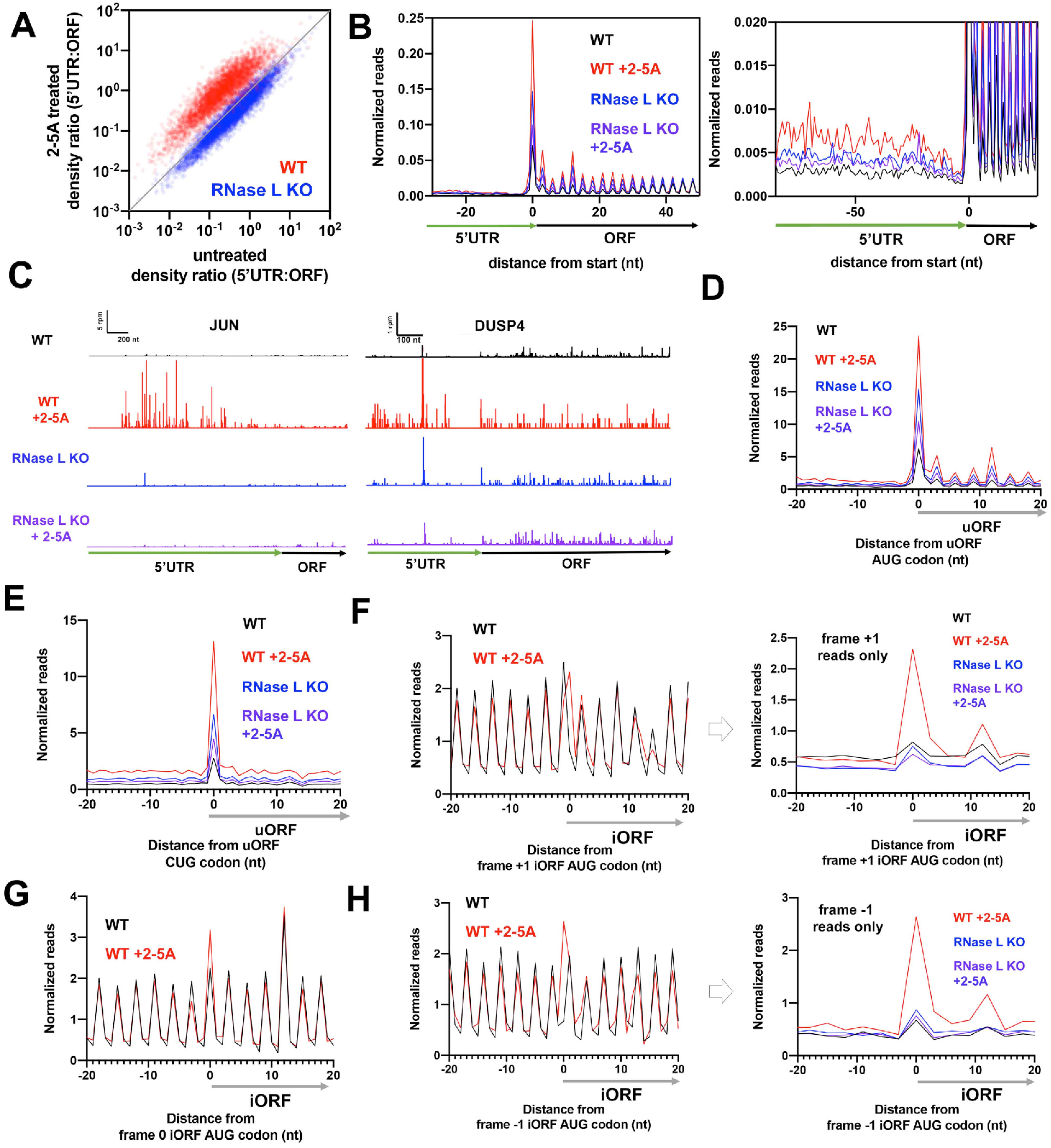
Translation of uORFs and iORFs is enhanced relative to coding sequences during RNase L activation. See also Supplemental Figure 3. **A** Comparison of 5’UTR:ORF ribosome footprint density ratios in 2-5 A treated versus untreated conditions is shown for WT and RNase L KO A549 cells. Each dot represents one gene model. Genes plotted above the diagonal have increased relative 5’UTR translation in treated in contrast to the untreated sample. **B** Normalized ribosome footprint occupancy (metagene plot) around the start codon of main ORFs reveals increased ribosome footprint levels in 5’UTRs. **C** Example gene models showing increased 5’UTR footprints in 2-5 A treated WT cells relative to untreated. Data from RNase L KO cells also shown as a control. Note that y-axis is truncated causing the highest peaks in 3’UTRs to be higher than shown for JUN and DUSP4. **D** Average ribosome footprints around AUG start codons in 5’UTRs show an increased peak at the start codon followed by footprints that exhibit 3-nt periodicity in 2-5 A treated WT cells, but not in RNase L KO cells. Ribosome footprints are normalized to main ORF footprint levels prior to averaging. **E** Average ribosome footprint levels as in D, but for CUG start codons, also show an increased peak. **F** Average ribosome footprints around AUG start codons in the +1 reading frame of coding sequences show an increased peak in 2-5 A treated WT cells (left panel). To further emphasize differences, we eliminate reads from the main ORF by plotting frame +1 reads only (right panel). **G** Average ribosome footprints around AUG start codons internal to and in the same frame as the main ORF of coding sequences show an increased peak in 2-5 A treated WT cells. **H** Average ribosome footprint levels as in F but for the −1 frame.

These increases were also evident in metagene plots where we averaged ribosome footprints from all genes that had been aligned by their canonical start codons (Figure 4B, Supplemental Figure 3D). We noted that the fold increase in 5’UTR ribosome occupancy was generally smaller as compared to 3’UTRs, mainly because 5’UTRs are normally translated at a moderate level, unlike 3’UTRs (Ingolia et al., 2014) (Figure 2B vs 4B, right-hand panels). Similar to the increase of ribosome protected footprints in 3’UTRs, we also noted these effects on individual genes (Figure 4C). We further investigated whether these trends were caused by active translation of individual uORFs by computing metagene plots aligned at uORF start codons. We found that overall uORF translation, as evidenced by increased peaks on start codons and overall higher footprint levels across the uORFs, was enhanced in 2-5A treated cells (Figure 4D-E and Supplemental Figure 3E and F). As with dORFs, 3 nt-periodicity downstream of AUG-initiated uORF start codons was evident, but less well-resolved for the case of non-canonical CUG codons (Figure 4D-E).

### RNase L activation increases internal ORF (iORF) translation

One hypothesis that could explain the increased translation of ORFs in UTR regions is that ribosomes are able to directly load onto mRNA fragments that are created by RNase L and begin translation at an available start codon. To test this prediction, we also asked whether alternative translation events take place within the coding sequences. To this end, we examined whether initiation of translation could take place at AUG codons found within the main ORF. Initiation of translation at such internal AUGs could result in one of two possible outcomes. If the AUG was in the same frame as the main ORF (frame 0), we would expect an N-terminally truncated protein product compared to that encoded by the full main ORF. If the AUG was in the −1 or +1 frame with respect to the main ORF, we would expect a ribosome to initiate and then terminate translation within an internal ORF (“iORF”) and generate a short peptide product. Either event (in-frame or out-of-frame initiation) should be detectable by the averaging (metagene) approach we previously used for uORFs and dORFs. However, any signal from these events will be confounded by translation of the main ORF. In the case of initiation in frame 0, we would expect a higher peak of ribosome footprints on the internal AUG codons due to the additional reads from that initiation event. In the case of initiation in frame +1 or −1, we would expect this peak to occur 1-nt away from the dominant 3-nt signal arising from the main ORF.

We therefore performed metagene analysis at internal AUG start codons (Fig 4F-H), similar to the approach we used for uORF and dORF metagene analysis. However, here we analyzed translation of these events in the different frames (frame 0, −1 and +1) individually to clearly distinguish footprints deriving from internal initiation events from the dominant main ORF footprints. We observed increased ribosome footprints, on average, at internal start codons in all frames in 2-5A treated cells (Figure 4F-H). In the case of out-of-frame iORF initiation, we further analyzed translation downstream of the initiation event by plotting only footprint reads in the frame corresponding to the iORF (−1 or +1). This eliminates the signal from main ORF translation and clearly shows continued elongation past the initiation event (Figure 4F and H, right panels). However, we could not analyze periodicity since this iORF analysis is limited single frame (+1 or −1). This analysis was consistent across replicates (Supplemental Figure 3G-I). As an internal control, we also analyzed out-of-frame reads in one case (−1 reads for +1 iORFs) and confirmed that we only observed footprints in the expected frame (Supplemental Figure 3I, note peak is only present in +1 frame). This analysis, combined with our previous analysis of uORFs and dORFs, comprehensively examines translation across the entire mRNA and shows that relative increases in alternative ORF (altORF) translation during RNase L activation are not limited to any particular region.

### The dsRNA mimic, poly I:C, induces altORF translation

While we found that altORFs in all mRNA regions were translated upon the specific activation of RNase L by 2-5A, we asked whether the effect would also manifest during broad activation of the innate immune response by the dsRNA mimic, poly I:C. We expect that during poly I:C treatment, RNase L would be activated by naturally synthesized 2-5A because poly I:C activates the OAS enzymes and, to varying degrees, other dsRNA sensors, such as RIG-I, MDA5, TLR3, and PKR (Alexopoulou et al., 2001; Hartmann, 2017; Kato et al., 2006). However, the activation of all of these pathways could mask the effects of RNase L activation. We therefore transfected A549 cells with poly I:C (0.25 μg/ml, 4.5 h), and performed ribosome profiling. We found that the effect of poly I:C on altORF translation was similar to 2-5A (Figure 5A-F, Supplemental Figure 4A-F). We observed increased footprints in dORFs in 3’UTRs (5A-D), uORFs in 5’UTRs (5E), and iORFs in coding sequences (5F) upon poly I:C transfection. Importantly, we observed a strong correlation in relative 3’UTR footprint levels observed between 2-5A and poly I:C treatment (Pearson’s R^2^=0.60 and 0.85 (replicate 1 and 2), Supplemental Figure 4B). We also detected a large increase in average ribosome footprint peaks on AUG codons of dORFs when cells were treated with poly I:C (Figure 5D and Supplemental Figure 4D). All of these effects of poly I:C treatment were substantially diminished in the RNase L KO cells (Figure 5A-D). The similarity in the effects suggests that translation of non-canonical regions occurs when RNase L is activated via naturally produced 2-5A from broad activation of the antiviral response by double-stranded RNAs. It should be noted that poly I:C treatment did result in slightly elevated 3’UTR ribosome footprints on some genes in a RNase L KO cell line (Supplemental Figure 4A). However, this increase across replicates was generally weak (Supplemental Figure 4C), the footprint pattern did not match that observed for 2-5A treated cells (Supplemental Figure 4G), and, in particular, peaks were not evident on AUG codons (Figures 5D, Supplemental Figure 4D)). This suggests that poly I:C may affect 3’UTR translation via a weak, alternative mechanism that is independent of RNase L.

**Figure 5.**
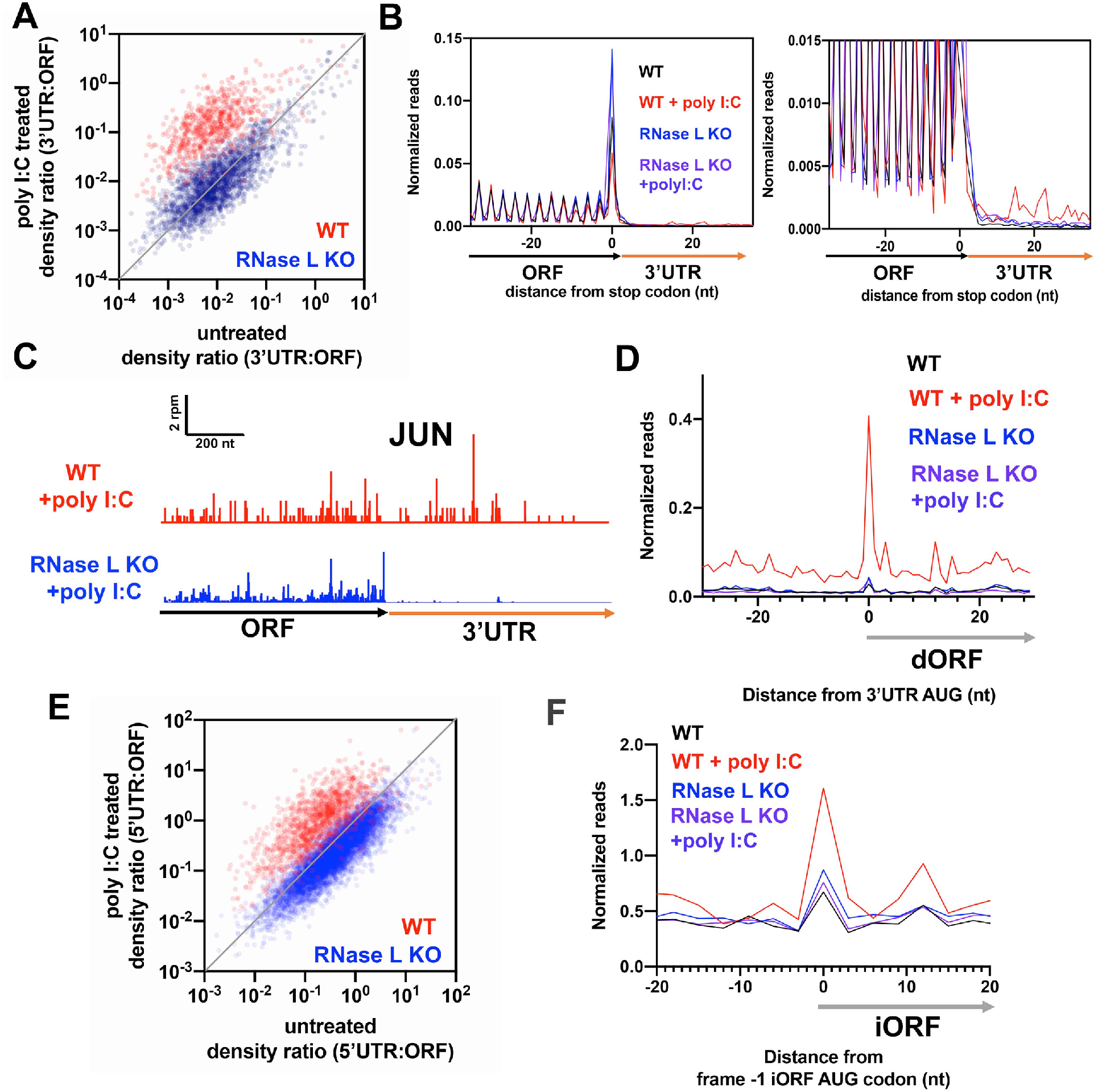
dsRNA enhances altORF translation. See also Supplemental Figure 4. **A** Comparison of 3’UTR:ORF ribosome footprint density ratios in 0.25 μg/ml poly I:C treated as compared to untreated conditions for WT and RNase L KO A549 cells. Each dot represents one gene model. Genes plotted above the diagonal have increased relative 3’UTR translation in the treated vs untreated sample. All plots include data from genes that passed threshold in both datasets (at least 2 raw reads in both the 3’UTRs in the main ORFs). **B** Normalized ribosome footprint occupancy (metagene) around the stop codon of main ORFs reveals increased ribosome footprint levels in the 3’UTRs in poly I:C treated cells as compared to untreated WT and RNase KO cells. **C** *JUN* gene model showing increased 3’UTR ribosome footprints in 2-5 A treated WT vs RNase L KO A549 cells. **D** Average ribosome footprints around AUG start codons in 3’UTRs show an increased peak in at the start codon in poly I:C treated WT cells, but not in RNase L KO cells. **E** Comparison of 5’UTR:ORF ribosome footprint density ratios in poly I:C treated vs untreated conditions for WT and RNase KO cells. All plots include data from genes that passed threshold in both datasets (at least 2 raw reads in both the 3’UTRs in the main ORFs). **F** Average ribosome footprints around AUG start codons in the −1 reading frame of coding sequences show an increased peak in poly I:C treated WT cells (left panel). To further emphasize differences, we plot frame −1 reads only (right panel), similar to Figure 4G.

### Catalytic activity of RNase L is required for alternative translation events

Having established that activation of RNase L increases the relative translation of altORFs, we further investigated the mechanism of this phenomenon by testing whether the catalytic activity of RNase L was required. By cleaving mRNAs, RNase L generates 5’ ends that could be used by 40S ribosomal subunits for initiation on mRNA fragments. Alternatively, it is plausible that RNase L could promote translation of altORFs via interactions with translation factors independent of RNA cleavage by, for example, increasing usage of a leaderless (5’ end independent) translation initiation mechanism (Andreev et al., 2006) (see Discussion).

First, we checked whether the relative increase in ribosome footprints in 3’UTRs is correlated with levels of RNA cleavage by RNase L. We reduced RNA cleavage by decreasing the 2-5A treatment duration from 4.5 to 2.25 hours and evaluated the result with the rRNA cleavage assay (Supplemental Figure 5A, lane 6 vs Figure 1B, lane 2). We found that 3’UTR footprints increased to a lesser extent compared to that of the 4.5 h treatment (Supplemental Figure 5B). Consistent with this, reducing 2-5A from 1 μM to 0.1 μM did not yield any detectable rRNA cleavage and did not increase the 3’UTR:ORF density ratio in ribosome profiling experiments (Supplemental Figures 5A, lane 2 and 5C). These data show that rRNA cleavage by RNase L is a good predictor of the relative increase in altORF translation and is consistent with a model where the cleavage activity of RNase L plays a role in translation of altORFs.

Next, we transiently expressed WT or H672N (catalytic mutant) RNase L in the RNase L KO cell line and examined changes in ribosome distribution by ribosome profiling (Figure 6A, Supplement Figure 5D). The H672N RNase L was shown to be deficient in cleavage activity due to the mutated catalytic histidine in the active site. Despite this mutation, it was also shown to maintain the ability to dimerize in the presence of 2-5A and bind RNA (Han et al., 2014). Therefore, we expect that this mutant should also maintain the protein-protein and protein-RNA interactions of wild-type RNase L. We confirmed the cleavage activity of transiently-expressed wild type RNase L by activating it with 1 μM 2-5A or 0.25 μg/ml poly I:C and performing rRNA cleavage assays (Figure 6B). In contrast, transiently-expressed mutant RNase L could not cleave rRNAs during 2-5A and poly I:C treatment (Figure 6B). As anticipated, we found that the 3’UTR:ORF ribosome footprint density ratio was higher when the transiently-expressed wild type RNase L was activated (2-5A or poly I:C) as compared to when the catalytic mutant was expressed (Figure 6C). We noted, however, that the extent of the effect was somewhat less than that with endogenous RNase L, likely due to the inherent inefficiency of the transient expression. This demonstrates that the relative increase in altORF translation is dependent on the catalytic activity of RNase L. In further support, local 3’UTR ribosome footprint patterns (Figure 6D) and the characteristic 3’UTR AUG peaks in average footprint data (Figure 6E and F) were present when wild type, but not mutant H672N, RNase L was transiently expressed in RNase L KO cells. We also note that poly I:C treatment on H672N expressing cells conferred increased footprints on some individual 3’UTRs (Figure 6D, ACTG1 data, row 6), as we noted in RNase L KO cells previously (Figure S4H). As in RNase L KO cells (Figure S4H), these footprints did not exhibit the same pattern of occupancy as in poly I:C-treated cells transfected with WT RNase L (Fig. 6D, rows 3 and 6). In particular, these footprints were not increased on start codons (Figure 6F).

**Figure 6.**
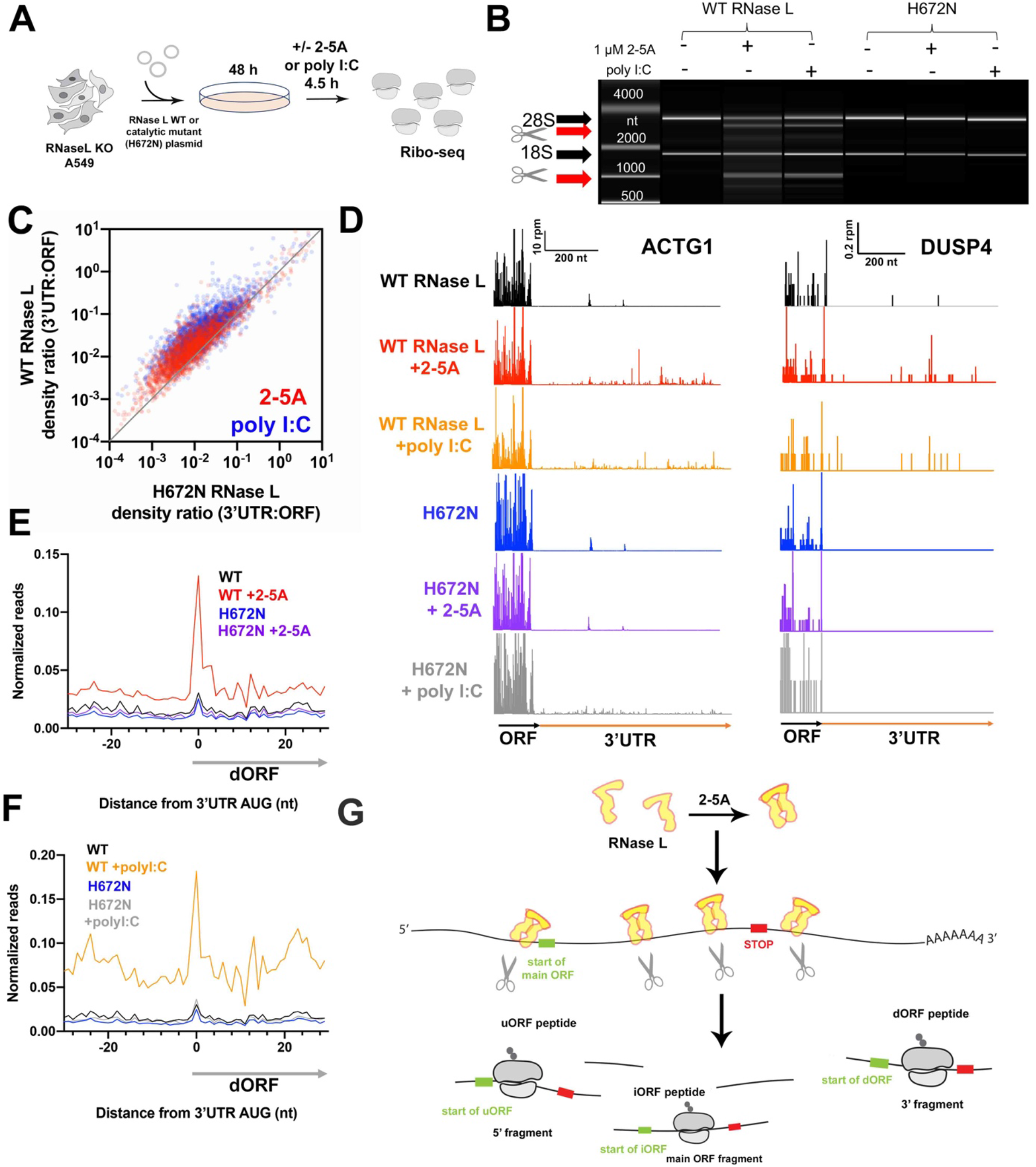
RNA cleavage activity of RNase L is required for altORF translation. See also Supplemental Figure 5. **A** Schematic of experiments. First, full length WT or H672N RNase L was introduced on a plasmid to A549 RNase L KO cells by electroporation. Then cells were then incubated for 48 hours and treated with 2-5 A or poly I:C for 4.5 hours. Cells were then harvested for Ribo-seq experiments. **B** BioAnalyzer-based rRNA cleavage assay showing activation when cells expressing WT RNase L are transfected with 2-5 A or poly I:C, but not the mutant H672N RNase L. **C** Comparison of 3’UTR:ORF ribosome footprint density ratios in 2-5 A or poly I:C treated conditions for RNase L KO cells that express WT vs H672N mutant RNase L. Each dot represents one gene model. Genes plotted above the diagonal have increased relative 3’UTR translation in the treated vs untreated sample. Plot includes data from genes that passed threshold in both datasets (at least 2 raw reads in both the 3’UTRs and in the main ORFs). **D** Footprints that map to a gene model of ACTG1 in RNase L KO A549 cells that express WT or H672N RNase L. Data show that treatment with 2-5 A or poly I:C increases relative 3’UTR levels of ribosome footprints. **E-F** Average ribosome footprints around AUG start codons in 3’UTRs show an increased peak at the start codon in 2-5 A (F) or poly I:C (G) treated cells that express WT, but not H672N, RNase L. Ribosome footprints are normalized to main ORF footprint levels prior to averaging. **G** Model to account for increased relative altORF translation during 2-5 AMD. First, RNase L dimerizes in the presence of 2-5 A and cleaves host mRNAs. Next, ribosomes load onto some fragments and translate encoded altORFs. The position of altORF start and stop codons are indicated on the translated fragments.

This inability of the catalytic mutant RNase L to induce alternative mRNA translation together with the correlation between rRNA cleavage and relative 3’UTR ribosome occupancy indicates that the observed increase in relative translation of altORFs depends on the catalytic activity of RNase L.

### Model of RNase L mediated altORF translation

Since we found that relative changes in altORF translation during RNase L activation are dependent on the cleavage activity of RNase L, we postulate that these effects rely on the presence of RNase L generated mRNA fragments. These fragments could potentially be translated if 40S ribosomal subunits can attach to the exposed 5’ ends and initiate translation at altORFs (Figure 6G). One prediction of this model is that these mRNA fragments should be stable enough to be identified by RNA sequencing methods. In support of this, it has been shown that rapid activation of RNase L leads to the accumulation of 3’UTR fragments (Rath et al., 2015). We investigated the nature of this phenomenon further on a global level by analyzing published data from a polyA^+^ RNA-seq experiment performed on poly I:C treated A549 cells (Rath et al., 2019). In this experiment, A549 cells were treated with 1 μM poly I:C for 4 hours, similar to the conditions that we employed for Ribo-seq. Much like our analysis of ribosome footprint analysis, we computed the ratio of 3’UTR:ORF polyA^+^ RNA-seq density and compared it between the untreated and poly I:C treated cells. We observed an enrichment in reads derived from 3’UTRs (Supplemental Figure 5F), indicating that stable 3’UTR fragments are present in the cell and could be translated as envisioned in Figure 6G. While the previous experiments did not address the status of fragments derived from 5’UTRs or coding sequences, we surmise that they are also likely present in the cytoplasm.

### Alternative ORFs are translated during RNase L mediated antiviral response

While we found that altORF translation occurs during activation of RNase L, it remains an open question if these non-canonical translation events also occur during viral infections. To answer this question, we took advantage of existing ribosome profiling data of viral infected cells. Vaccinia virus (VACV), which is part of the Poxvirus family, is known to be one of the most potent activators of the 2-5A mediated antiviral response (Rice et al., 1984). While this virus has a large dsDNA genome, late viral RNAs are thought to form dsRNA that activate the OAS-RNase L pathway (Rice et al., 1984). We analyzed publicly available Ribo-seq datasets from HeLa cells infected with VACV for variable durations (mock treated and VACV treated, 2h-8h) (Dai et al., 2017). We found that 3’UTR:ORF ribosome footprint density ratios increased in proportion to the length of treatment, as compared to a mock treated control, with 8 hours treatment having the greatest effect (Fig 7A, Supplemental Figure 6A). The magnitude of the increase was comparable to what we observed in A549 cells treated with 2-5A (Figure 2A). The effects were also detectable, on average, in a metagene plot computed from reads mapping near the stop codon and at the individual gene level (Figure 7B and C). Analysis of the dORF metagene plot for AUG codons also indicated specific translation of dORFs, as indicated by the peak at the start codon (Figure 7D). We also noted increases in uORF translation in 5’UTRs (Figure 7E, Supplemental Figure 6B). These Ribo-seq experiments on cells infected with VACV therefore indicate that translation of altORFs during the antiviral response can occur and may play a functional role defense against certain classes of viruses.

**Figure 7.**
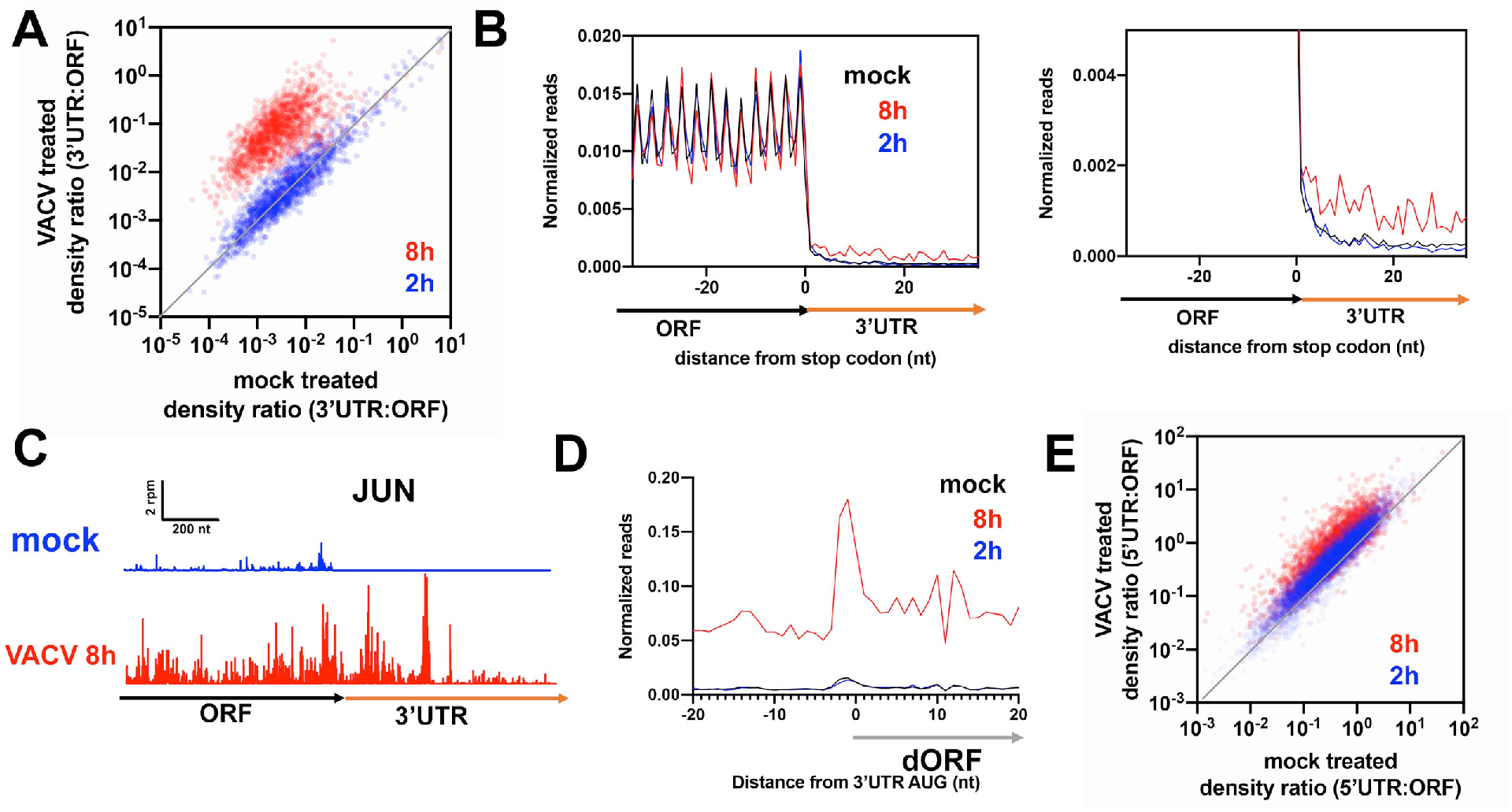
AltORFs are translated during vaccinia viral infection. **A** Comparison of 3’UTR:ORF density ratios in VACV vs mock treated (2 h, 8 h) HeLa cells. Each dot represents one gene. Genes plotted above the diagonal have increased relative 3’UTR translation in the treated vs untreated sample. **B** Normalized ribosome footprint occupancy (metagene plot) around the stop codon of main ORFs reveals increased ribosome footprint levels in the 3’UTRs in VACV treated (8 h) cells. **C** *JUN* gene model showing increased 3’UTR reads in VACV treated cells. **D** Average ribosome footprints around AUG start codons in 3’UTRs show an increased peak at the start codon in 8h VACV treated cells, but not in mock treated or 2 h treated cells. **E** Comparison of 5’UTR:ORF density ratios of transcripts in VACV vs mock treated (2h, 8 h) HeLa cells. Genes plotted above the diagonal have increased relative 5’UTR translation in the treated vs untreated sample.

## DISCUSSION

Activation of RNase L is known to be important in the defense against viral infections, yet it remains mostly unclear what effects mRNA degradation caused by this activation have on the host. We investigated how altered translation of the modified transcriptome might contribute to this function by performing ribosome profiling experiments on RNase L activated cells. We found that ORFs within non-coding mRNA sequences (altORFs, including uORFs, iORFs and dORFs) become translated in higher proportion in RNase L activated cells compared to resting cells. Notably, this altORF translation was dependent on the cleavage activity of RNase L. This finding disfavors models where translation termination and recycling are directly modulated and instead points to the importance of mRNA fragmentation in explaining this phenomenon. One possible model, based on data presented here and elsewhere (Rath et al., 2015; Rath et al., 2019), is that the RNA fragments cleaved by RNase L can be translated prior to being fully degraded (Figure 6G). In support of this model, RNase L dependent mRNA fragments were previously found in S10 HeLa cell lysates and T45D cells by poly A^+^ RNA-seq analysis upon 2-5A treatment (Rath et al., 2015). In addition, we showed that data from others (Rath et al., 2019) demonstrate that these mRNA fragments are stable enough to be detected in cells.

These observations raise the question of how these fragments are stabilized to avoid rapid degradation. One possibility is that components of the major RNA degradation pathways, such as Xrn1 (5’ and 3’ exonuclease) or subunits of the exosome (3’ to 5’ exonuclease), are inhibited because they become overwhelmed by the sheer number of cleaved mRNA substrates. Xrn1 was already shown to be an important regulator of the innate immune response due to its ability to degrade viral dsRNA fragments and thereby tune the host’s ability to detect the virus via other dsRNA sensors (Burgess and Mohr, 2015; Liu and Moss, 2016). Thus, inhibition of Xrn1 could play a role in setting the stability of both host and viral RNA fragments in the cell during RNase L activation. Furthermore, the RNase L cleavage reaction produces RNA termini, a 5’-OH on the 3’ cleavage fragment and a 2’-3’-cyclic-phosphate on the 5’ cleavage fragment (Cooper et al., 2014; Wreschner et al., 1981), that are likely to afford some resistance to degradation because they are not ideal substrates for Xrn1 and the exosome, respectively (Pellegrini et al., 2008; Shigematsu et al., 2018; Sporn et al., 1969).

Another question raised by this model is how 40S ribosomal subunits are recruited to these fragments in order to initiate translation. Canonical translation starts with the loading of the 40S subunit onto the 5’ end of the mRNA. This process is enhanced by interactions with initiation factors, particularly eIF4E that directly binds the the 5’ 7-methylguanylate (m7G) cap that is attached to mRNAs in the nucleus (Pelletier and Sonenberg, 2019; Yue et al., 1997). In the case of RNase L activation, the absence of a cap on all fragments (except the 5’-most fragment) would be expected to lower the efficiency of ribosome recruitment. However, many examples of cytoplasmic mRNA “recapping” have been demonstrated (Otsuka et al., 2009; Trotman and Schoenberg, 2019) and could account for why these fragments are translated. This sort of recapping process was shown to stabilize endonucleolytic decay intermediates that are generated during NMD in a β-thalassemic mouse model (Lim et al., 1989; Otsuka et al., 2009). On the other hand, translation of these fragments could also occur in the absence of a 5’ cap by alternative mechanisms, such as use of A-rich sequence elements that have been shown to promote initiation (Jia et al., 2020) or leaderless translation (Andreev et al., 2006). During leaderless translation, 80S ribosomes loaded with initiator Met-tRNA are believed to bind directly to start codons without need of initiation factors or a 5’ cap. Additionally, it is conceivable that there would be an overabundance of free ribosomes during RNase L activation because most mRNAs are degraded, freeing up ribosomes that would normally be engaged in translating conventional capped mRNAs. In this case where “ribosome homeostasis” is perturbed, it is predicted that low-efficiency, cap-independent recruitment of 40S ribosomal subunits would be enhanced simply due to mass action (Mills and Green, 2017).

While the observation of altORF translation can be explained by a model where mRNA fragments that have escaped degradation are translated (Figure 6G), other possibilities can be considered. For example, it is conceivable that RNase L cleavage might damage ribosomal RNA (rRNA) to an extent that alters the fidelity of translation initiation in a way that permits these damaged ribosomes to translate regions outside of the main ORF. It is also possible that changes in the abundance of initiation factors during the broad reprogramming of the transcriptome, or activation of stress pathways as a result of mRNA loss, could facilitate translation of altORFs. It was also recently proposed that dORFs in the 3’UTR can directly recruit ribosomes and lead to initiation (Wu et al., 2020b). Thus, it is possible that RNase L activation could also modulate this process.

Our observation also raises the question of whether translation of these altORFs affects the innate immune response and offers any benefit to the host in clearing virus-infected cells. We predict that some of the RNase L cleaved mRNA fragments contain a start codon, where translation can be initiated, but lack an in-frame stop codon due to cleavage at the 3’ end. Therefore, we speculate that ribosomes translating such fragments will become trapped at the 3’ end of the RNA and require rescue by ribosome rescue factors, such as PELO (Guydosh et al., 2017). This highlights a potential role of ribosome rescue factors during RNase L activation in maintaining a pool of free ribosomes. Prolonged activation of RNase L could overwhelm the ribosome rescue system and lead to ribosome queues that are, in turn, recognized by sensors of ribosome stalling, such as ZAKα, that could trigger apoptosis mediated by JNK (Vind et al., 2020; Wu et al., 2020a). Such a model could account for the observation that JNK is critical to RNase-L mediated apoptosis (Li et al., 2004), a process thought to be beneficial for eliminating cells infected by viruses.

The peptide products that are produced by altORF translation during RNase L activation could also have functional roles. It is known that translation outside annotated sequences can have important functional outcomes (Ingolia et al., 2014) and, according to the “immunogenic peptide hypothesis,” altORFs can be a source of cryptic (unannotated) peptides presented by Human Leukocyte Antigen I (HLA-I) molecules (Boon and Van Pel, 1989; Yewdell, 2011). If such peptides are seen as “non-self” by the adaptive immune system, they could aid in the clearance of viruses. In support of this idea, prior work demonstrated that cryptic peptides that are encoded by uORFs and dORFs can be effectively loaded onto HLA-Is (Chen et al., 2020; Schwab et al., 2003) and, in B-cells, 7.5% and 24.6% of the HLA-I bound peptidome originated from 3’UTRs and 5’UTRs, respectively (Laumont et al., 2016). It is therefore conceivable that peptides generated from translation of RNase L cleavage fragments could also be presented on HLA-I and benefit the host, acting as additional danger signals. However, given the short time frame of RNase L activation, detection of cryptic peptides in RNase L activated cells has proven to be difficult (data not shown). Thus, methodology development is needed to assess whether altORF-derived peptides are stable and functional.

In addition, it is possible that translation of the mRNA decay intermediates generated by RNase L could serve other roles. For example, they could sequester ribosomes and thereby prevent translation of viral transcripts. On the other hand, translation of fragments could be detrimental to the host since ribosomes melt mRNA secondary structure (Mizrahi et al., 2018) and this structure in the fragments was suggested to be important for activating other dsRNA sensors that trigger interferon production and assembly of stress granules (Burke et al., 2019; Malathi et al., 2007; Manivannan et al., 2020).

Finally, it is notable that the relative levels of altORF translation in cells infected with vaccinia virus (VACV) mimicked that in RNase L activated cells. This suggests that altORF translation is triggered by at least some viral infections. However, since many viruses are known to suppress activation of RNase L, the potential role of altORF translation in the clearance of the viruses is likely to vary. Therefore, more targeted studies are needed to investigate the role of RNase L in altORF translation in the context of viruses. Nevertheless, our findings reveal that widespread translation outside of coding sequences is a component of the innate immune response and add to the growing body of examples of alternative translation events. We believe further study of this phenomenon has the potential to identify new mechanistic targets for therapeutic intervention.

## ACKNOWLEDGEMENTS

We thank Dr. Robert Hogg and Dr. Nazmul Haque for valuable discussions and providing dual luciferase and RNase L plasmids. The A549 WT and RNase L KO cell lines were a kind present from Dr. Bernard Moss. OAS1(p42) plasmid was a generous gift of Dr. Pia Moller Martensen. We also thank Dr. Behdad Afzali and Dr. Daniel Chauss for their assistance with electroporation experiments. We are thankful to Dr. Bret Hassel for providing 2-5A for initial experiments. We are also grateful for valuable discussions and/or feedback on the manuscript from Dr. Alan Hinnebusch, Dr. Jon Lorsch, Dr. Tom Dever and Dr. Bret Hassel. This work was funded by the Intramural Research Program of the NIH, the National Institute of Diabetes and Digestive and Kidney Diseases (NIDDK) (DK075132 to N.R.G.) and the Postdoctoral Research Associate Training Program (PRAT) at the National Institute of General Medical Sciences (NIGMS) (1FI2GM128743 to A.K.).

## AUTHOR CONTRIBUTIONS

A.K. designed and performed experiments, analyzed the data and wrote the paper. G.D.J. contributed to preliminary data and analysis and performed ribosome profiling of Hap1 cells. A.V.D. performed experiments for preliminary data. N.R.G. designed experiments, developed software, analyzed the data and wrote the paper.

## STAR METHODS

## CONTACT FOR REAGENT AND RESOURCE SHARING

Further information and requests for resources and reagents should be directed to and will be fulfilled by the Lead Contact, Nicholas Guydosh.

## EXPERIMENTAL MODEL AND SUBJECT DETAILS

### Tissue Culture Cells

Human cell lines Hap1, HeLa and A549 were cultured in DMEM (Hap1, HeLa) or RPMI (A549) complemented with 10% Fetal Bovine Serum (FBS). A549 cells were tested and negative for Mycoplasma contamination throughout the study. Mycoplasma testing was performed using eMyco Valid Mycoplasma PCR detection kit (BioLink). Cells were incubated at 37°C in the presence of 5% CO_2_.

## METHOD DETAILS

### Synthesis and purification of 2-5A

2-5A was synthesized enzymatically *in vitro* by recombinant human OAS1. First, recombinant OAS1 (p42) containing an N-terminal His tag was expressed in BL21(DE3) *E.coli* as described before (Poulsen et al., 2015) or with an alternative protocol using autoinduction media (Studier, 2005). Then cells were collected by centrifugation at 7000 g for 15 min at 4°C. Then cells were lysed in B-per protein extraction reagent 4 ml/ gram bacterial pellet in the presence of cOMPLETE mini protease inhibitor cocktail (Roche) for 15 minutes at room temperature. Next, bacterial lysate was cleared by centrifugation at 34,000g for 1 hour at 4°C. The supernatant was filtered (45 μm pore size filter, Millipore) and loaded onto a HisTrap (GE Healthcare) nickel column. The column was then washed with wash buffer (20 mM Hepes (pH 7.5), 300 mM NaCl, 10% (vol/vol) glycerol, 1 mM TCEP and 50 mM imidazole) and OAS1 was gradient eluted with 500 mM imidazole. Protein fractions were then evaluated by SDS-PAGE and Coomassie staining. Then OAS1 containing fractions were pooled, concentrated and buffer exchanged (Zeba Spin desalting Column, 7 K MWCO, Thermo Scientific) in storage buffer (20 mM Hepes (pH 7.5), 300 mM NaCl, 10% (vol/vol) glycerol and1 mM TCEP). Final protein preparation was stored in storage buffer at −80°C. Concentration of OAS1 was determined by using NanoDrop Spectrophotometer (Thermo Fischer Scientific) (Mw=41.5 g/mol and ɛ= 65,485 M^−1^cm^−1^).

To produce 2-5A, 2 μM purified OAS1 was incubated with 1.25 OD_260_ poly I:C in the presence of 10 mM ATP, 20 mM Hepes (pH 7.5), 50 mM NaCl, 30 mM MgCl_2_, 10% (vol/vol) glycerol, 4 mM DTT at 30°C for 2 hours. To stop the reaction, samples were incubated at 85 °C for 15 minutes. Then the samples were filtered (22 μm pore size Millipore filter) and the different 2-5A species were separated on a 16/10 MonoQ column as described before (Poulsen et al., 2015). The same 2-5A fractions from several run was pooled and run again on 16/10 MonoQ column to achieve higher concentrations. 2-5A concertation was estimated by NanoDrop Spectrophotometer at 259 nm. Then yielded 2-5A was aliquoted and stored at −80°C.

### Ribosome profiling

Cells used in experiments were grown until they reached near confluency (70-90% confluent) in a T75 flask. Then cells were transfected with the pre-incubated mixture of 0.1−10 μM 2-5A or 0.25 μg/ml poly I:C and Lipofectamine 3000 (58 μl/T75 flask) in Opti-Mem serum free media as recommended by the manufacturer. After incubation (2-4.5 h) cells were washed with ice-cold PBS and then flash frozen in liquid N_2_. 400-600 μl of lysis buffer (20 mM Tris-HCl pH 7.5, 150 mM NaCl, 5 mm MgCl_2_, 1 mM DTT, 100 μg/ml cycloheximide, 1% Triton X−100, 0.05 U Turbo DNase) was added to the frozen cells and those were thawed on ice. Cells were scraped from the bottom of the flasks and lysates were transferred to Eppendorf tubes and incubated on ice for additional 10 minutes before passing through 25 Gauge needles ten times. Lysates were clarified by centrifugation for 10 minutes at 21,000 g at 4°C. Supernatants were flash frozen in liquid N_2_ and stored at −80°C before proceeding to further steps. All other steps and construction of the Illumina sequencing libraries were carried out as described before (McGlincy and Ingolia, 2017). rRNAs were removed by Illumina Ribo-Zero rRNA removal kit. Quality of the libraries was assessed by Bioanalyzer 2100 (Agilent) using the High Sensitivity DNA kit (Agilent). Sequencing was performed on an Illumina HiSeq3000 at the NHLBI or NIDDK DNA Sequencing and Genomics Core.

Ribo-seq experiments with transiently transfected A549 RNAse L KO cells were carried out similarly to the above. Wild type and catalytic mutant H672N RNase L plasmids cDNAs (the backbone is pcDNA5-FRT-TO with CMV promoter, Invitrogen) used in this study were a generous gift from Dr. Robert Hogg. Originally, wild type RNase L was cloned from HEK293 cells and H672N mutation was introduced by quick change mutagenesis. The cDNA sequences were identical to that of the reference sequence encoding the full-length RNase L. For electroporation experiments confluent T75s (~ 6-8*10^6^ cells) were trypsinized and resuspended in MaxCyte electroporation buffer and electroporated in 3XOC25 cuvettes (MaxCyte) in the presence of 150 μg/ml pcDNA5-RNAse L wild type or H672N transfection quality plasmid with standard settings for A549 in the MaxCyte ATX electroporator. Cells were divided and plated on three T75 flasks and incubated for 48 h until cells reached 70% confluency before 2-5A or poly I:C treatment. All other steps were carried out as described above.

### Computational analysis

#### Read processing

The fastq files were de-barcoded by the core facility and reads of 25-34 nucleotides were sorted by internal 5 nucleotide sample barcode and trimmed of linkers by Cutadapt. Then contaminating tRNAs and remaining rRNAs were filtered out by Bowtie1 allowing two mismatches in -v mode. Additionally, we used the -y option to increase sensitivity of the alignment. We created the noncoding RNA fasta file by downloading rRNA sequences from the SILVA project (release 128) (Quast et al., 2013) and tRNA sequences from GtRNAdb (H. sapiens release 16) (Chan and Lowe, 2009). After the filtering out the non-coding RNAs, an in-house python script (dedup) was used to remove PCR duplicates by comparing the 7 nucleotide unique molecular identifiers (UMI). Then, the resulting fastq files were quality checked by FastQC. In particular, we found that the majority of the ribosome protected footprints were 28-29 nucleotides long. Then UMIs were trimmed by Cutadapt and reads were aligned to the human transcriptome. The hg38 genome and annotations originated at UCSC on August 14, 2015 and were downloaded from the Illumina iGenomes project. Tophat 2.1 and bowtie.1.2 aligner were used and allowed up to 2 mismatches (-N 2). The -T, -g 1, and --no-novel-juncs options were used as further inputs to allow a single mapping per read to annotated transcriptome only. The resulting SAM files containing aligned reads were sorted and indexed using Samtools for downstream analysis. In order to view reads in a genome browser, wig files were generated by using the make_wiggle function of the Plastid suite. We note that we used 5’-end assignment of footprints throughout the study since it offers a higher fraction of reads mapping to a single reading frame (reading frame of the main ORF) than 3’-end assignment or coverage. Initial read processing, bowtie alignments to ncRNA, and tophat alignments were performed on the NIH Biowulf cluster.

#### Analysis of data

##### Calculation of UTR:ORF density ratios

We quantitated the number of reads mapping to defined sub-regions (CDS, 5’UTR, and 3’UTR) by using the Plastid suite (Dunn and Weissman, 2016) and *cs* function. First, we used the *generate* mode to preprocess the annotation file used above and in the downstream analysis. Then we obtained counts of aligned reads in these regions with *count* mode using options for 5’ end alignment and offset of 13 nucleotides (corresponds to center of P site). Raw read values were normalized to the total number of reads that mapped in the alignment. We then computed the density (in units of rpkm) in each region by dividing the number of counts by the region’s length. Then the ratio of UTR and main ORF raw reads (densities) were calculated. Genes were excluded where there were less than 5 raw reads in both the UTR and ORF unless otherwise indicated.

##### Metagene analysis

We created average ribosome footprint density (metagene) plots by using Plastid’s *metagene* function. First, the *generate* mode was used to define a window of 400 nucleotides around the start and the stop sites that was free of alternative splicing. Then we used *count* mode to calculate the mean ribosome occupancy across genes. Metagene plots were normalized so that the contribution from each gene was equally weighted according to the density within its main ORF, specified by positions 10 to 100 and −100 to −20 from start and stop codon plots, respectively. Data traces are shifted so that peaks coincide with start or stop codons (approximate P site or A site, respectively). Note that peak 12-nt past start codon occurs because all reads at this site have ATG at their 5’ end and this amplifies with high efficiency during library creation.

##### Differential translation analysis

To determine changes in ribosome footprint distribution in RNase L activated cells we analyzed two complete and independent Ribo-seq datasets (replicate 1 and 2). Analysis were carried out by using DTEG.R as described (Chothani et al., 2019), that builds on the widely used differential expression analysis software DEseq2, however DTEG.R is able to take other factors, such as batch effect, into account. Data input for DTEG.R came from the Plastid cs function, where raw read count values were used. Gene ontology analysis was performed by GOrilla webserver (Eden et al., 2009) where enrichment of genes with the largest, significant increase (>4 fold) in ribosome occupancy were compared to the background set of genes (two unranked list of genes and function options).

#### Reduced transcriptome analysis

For more detailed analysis, we implemented a more stringent transcriptome-only analysis approach with a single transcript isoform per gene to enable higher precision mapping and eliminate spurious reads. In detail, we took reads that had been digitally subtracted for noncoding RNA and aligning them to a transcriptome with bowtie version 1.2.3 using the parameters -v 1 (1 mismatch allowed) and -y. The transcriptome, RefSeq Select+MANE (ncbiRefSeqSelect), was downloaded from UCSC on April 14, 2020 and used for alignment after removal of duplicates on alt chromosomes. We then used custom Python3 scripts to perform specialized average (metagene) analysis at start and stop codons.

##### Metagene analysis for reduced transcriptome alignment

Metagene analysis (metagene_m) were performed in a few cases for special cases of long windows where the high-quality transcriptome annotation was required using the transcriptome alignment described above (Supplemental Figure 1D). Analysis was analogous to that performed by Plastid. Averages were computed around the window size of choice for each gene after normalizing to the total read density of the main ORF. Genes with <5 rpkm read density were not included and traces shifted to correspond to approximate A site.

##### altORF metagene analysis

Averages (posavg_m) around start codons in UTR regions (Figures 2–7) were computed by normalizing reads in a window of 30 nt upstream and downstream of the codon of interest, after shifting them 12 nt to correspond the approximate P site, to the average read density in the main ORF. To minimize bad annotations, we excluded genes where main ORF footprint density measured <1 rpkm and cases where the average read density in the window was 10-fold greater than the main ORF. For averages of start codons within coding sequences (iORFs and N-terminal truncations), each start codon was equally weighted in the average according to the total reads that mapped within the window of interest. Note that the peak 12-nt past start codon occurs because all reads at this site have ATG at their 5’ end and this amplifies with high efficiency during library creation.

##### Catalog of dORF sites

Read density on all possible AUG-initiated dORFs was computed (dorflist) by summing reads mapping only in the frame of interest. These values were then used to compute the length of those dORFs whose relative expression most increased under 2-5A treatment (Figure 3E).

##### Gene models

Example ribosome profiling data on particular gene models was generated by using writegene2_m to show reads mapping to the reduced transcriptome alignment.

### rRNA cleavage assay

Total RNA was extracted from 100 μl of lysates prepared for ribosome profiling using Direct-zol mini kit (Zymo) or Trizol reagent (Invitrogen) according to the manufacturer’s protocol. The amount of total RNA was computed by absorbance at 260 nm measured by NanoDrop (Thermo Fischer Scientific) and then diluted to 50-500 ng/μl. Then RNA samples were run on Bioanalyzer 2100 (Agilent) using the RNA 6000 Nano kit (Agilent). Minute run-to-run shifts in band sizes are due to inherent limitations of the instrument.

### Dual luciferase 3’UTR translation assay

Dual Luciferase plasmids were previously generated by cloning Renilla Luciferase (rLuc) into the main ORF followed by a stop codon of the pT7CFE-CHis plasmid backbone (see in (Houck-Loomis et al., 2011)). To study 3’UTR translation Firefly Luciferase (fLuc) was inserted into the 3’UTR (3’UTR plasmid). These two reporter sequences were in the same frame and were separated with an 84 nt long linker. While fLuc did not contain its own AUG start codon, the linker encoded a non-canonical CUG start codon upstream of the cDNA of fLuc. As a control, the same cDNA sequence was used without the stop codon between the rLuc and fLuc sequences (control plasmid). The product of the control plasmid is expected to have both rLuc and fLuc activity and represent the case where every ribosome translates both genes. The assay was performed by incubating the plasmids with or without the trimer form of 2-5A in HeLa cell lysates at 30°C for 90 minutes using the 1-step Human Coupled IVT Kit (see details in key resources table). Then samples were then incubated on ice for 10 minutes to rapidly terminate the reaction. Luciferase activity was assayed by using a Dual-Glo Luciferase assay (Promega) to generate luminescence and measuring it with Berthold Centro LB 960 plate reader.

### Western blot analysis

The level of transient expression of RNase L for Ribo-seq experiments was evaluated by SDS-PAGE electrophoresis and immunoblotting. First, 10 μl of lysate was mixed with SDS Sample Buffer (Invitrogen) then run on 4-20% gradient Mini-PROTEAN Tris-HCl gel (BioRad). Proteins were transferred to a 0.2 μm PVDF membrane using the Trans-Blot Turbo System (BioRad) and membranes were blocked in Tris Buffered Saline, 0.1% Tween 20 (TBST) and 5% milk for 1 hour at room temperature. This was followed by incubation with primary antibodies overnight in TBST (Tris-buffered saline plus 0.1% tween) and 1% milk (RNase L antibody 1:10000, Histone H3 antibody 1:3000 dilution). After washing three times for 5 minutes in TBST, secondary antibodies were incubated with the PVDF membrane for 1 hour at room temperature (Goat anti-rabbit antibody, BioRad, 1:3000). Then, after additional washes (3 times, 5 minutes each) in TBST, the PVDF membrane were incubated with Clarity Western ECL substrate (BioRad) for 5 minutes and proteins were visualized by Amersham Imager 600.

## QUANTIFICATION AND STATISTICAL ANALYSIS

### Ribosome profiling

Ribo-seq experiments were repeated at least twice except electroporation experiments (Figure 5) where we used two different conditions (2-5A and poly I:C treatment) instead of replicates to support our conclusions. Each biological replicate dataset was analyzed individually as described above. Analysis and comparison of all datasets are provided in the main and in the supplement figures. We note that due to the variability in cell confluency and transfection efficiency of 2-5A and poly I:C we also see some differences between UTR:ORF ratios in biological replicates. However, all of the 2-5A and poly I:C treated WT A549 datasets exhibited increased UTR:ORF ratios when compared to their respective untreated dataset (transfection control). In the case of A549 RNase L KO datasets, while we found that nearly all of the 2-5A and poly I:C treated cells did not exhibit increased UTR:ORF ratios, we noted one exception (Supplemental Figure 3A). In this case, untreated cells (replicate 2) showed particularly low 5’UTR:ORF ratios (see boxplot in Supplemental Figure 3C), resulting in an apparent shift above the diagonal when 2-5A treated RNase L KO data were plotted against it (Supplemental Figure 3A). Since all 2-5A treated RNase L KO datasets exhibited lower 5’UTR:ORF ratios than 2-5A treated WT datasets (first 5 boxplots, Supplemental Figure 3C), the shift in this case can be attributed to inherent variability of 5’UTR occupancy in the data. All plots were created by using Prism 8.1.0 software or Igor Pro version 8. Correlation analysis (Pearson’s r of log-transformed data) was performed by using Prism 8.1.0 software. In box plots of UTR:ORF density ratios boxes represent the interquartile range (IQR) and horizontal line is the median. Whiskers show 1.5*IQR and notches give 1.58*IQR/√N.

### Dual luciferase 3’UTR translation assay

HeLa lysates without addition of plasmids were used to determine the background for rLuc and fLuc. In each experiment background values were subtracted from all data points. fLuc activities of 3’UTR and control plasmid were normalized based on the main ORF, rLuc activity. The amount of 3’UTR translation (fLuc activity from the 3’UTR plasmid) was then expressed as the percentage of fLuc activity of the control plasmid. Three independent experiments, each containing two technical replicates, and standard error of the mean (S.E.M) were plotted using Prism 8.1.0 software.

---

**S1.**
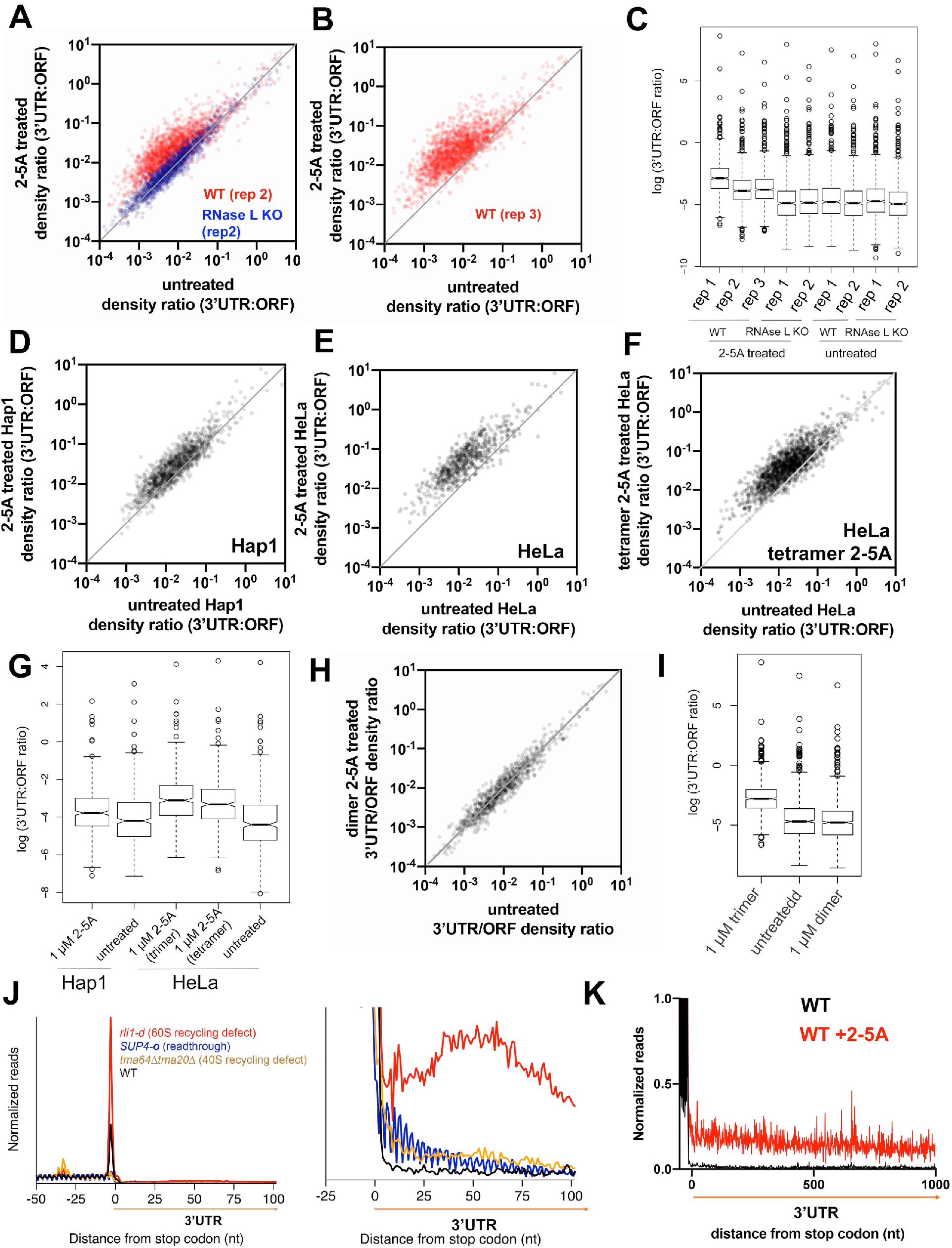
**Related to Figure 2. A-B** Comparison of 3’UTR:ORF ribosome density ratios of genes in trimer 2-5 A treated and untreated WT (2^nd^ and 3^rd^ replicates) and RNase L KO (2^nd^ replicate) cell lines. Genes plotted above the diagonal have increased 3’UTR:ORF ribosome footprint density ratios in treated sample as compared to that of untreated. **C** Box plot representation of data from Figure 2A and S1A and B. **D-F** Comparison of 3’UTR:ORF ribosome footprint density ratios in trimeric 2-5 A treated vs untreated conditions for Hap1 (D) or HeLa (E) cells, or tetrameric 2-5 A in HeLa cells (F). Each dot represents one gene model. Genes plotted above the diagonal have increased relative 3’UTR translation in the treated vs untreated sample. Data show increased 3’UTR translation occurs in multiple cell lines and that tetrameric 2-5 A also causes it. However, we note that Hap1 cells display lower density ratios as compared to Hela and A549 cells, likely due to the less transfection efficiency of this cell line. **G** Box plot representation of data shown in D-F, performed as in C. **H** Analysis of 3’UTR:ORF density ratios as in D-F, but for dimeric 2-5 A in A549 cells. Data reveal dimeric 2-5 A does not increase relative 3’UTR translation. **I** Box plot representation of H with data from trimeric 2-5 A also shown as reference. **J** Different mechanisms of 3’UTR translation in yeast (Young et al., 2015; Young et al., 2018). **K** Metagene analysis similar to Figure 2B, but limited to genes with long 3’UTRs. This shows minimal change in 3’UTR footprint levels in regions far downstream of the stop codon.

**S2.**
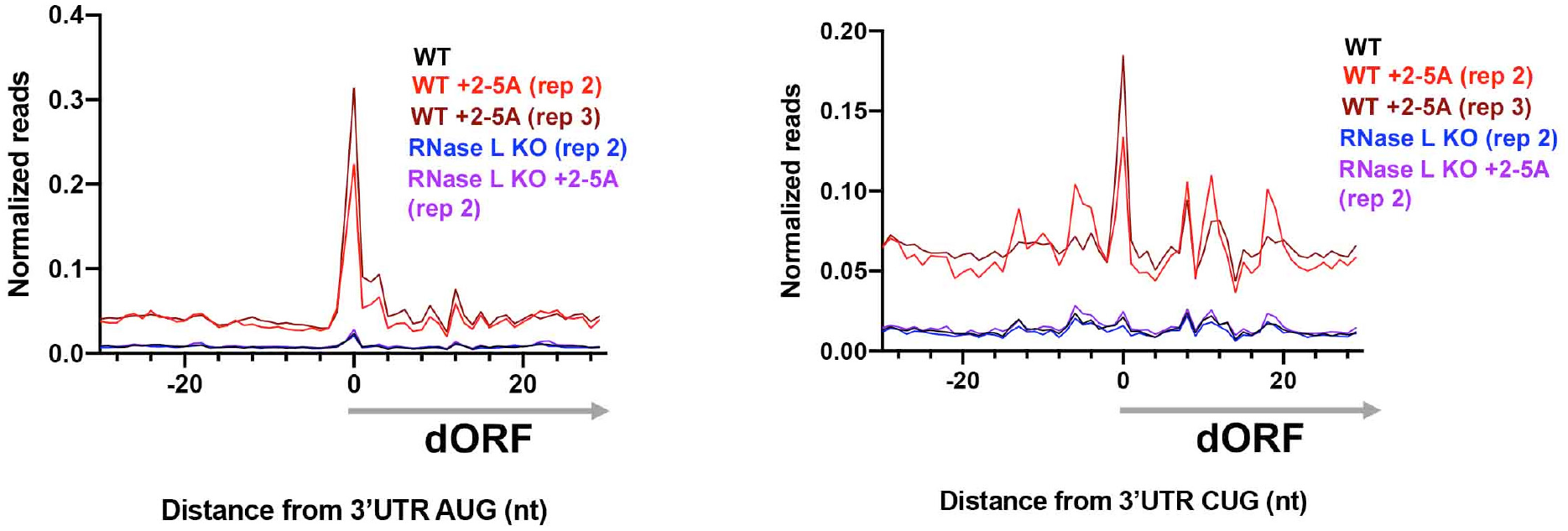
Replicate analysis confirms increased relative translation of dORFs after 2-5 A transfection. Related to Figure 3. 3’UTR metagene analysis of ribosome footprints showed increased AUG and CUG peaks in 2-5 A treated WT cells (2^nd^ and 3^rd^ replicates), but not in RNase L KO cells (2^nd^ replicate).

**S3.**
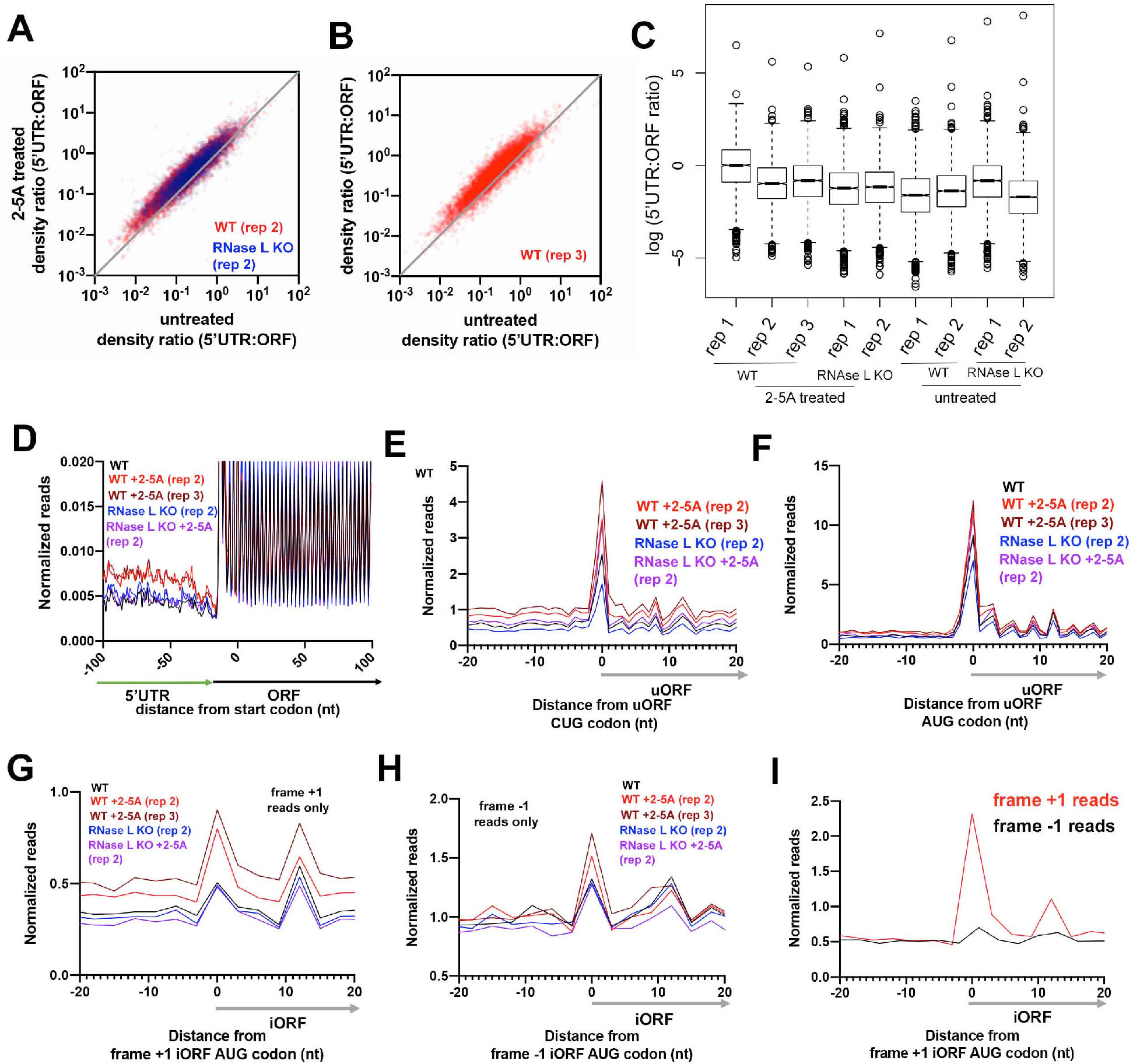
**Related to Figure 4.** Comparison of 5’UTR:ORF density ratios of transcripts in trimer 2-5A treated and untreated **(A-B)** WT (replicate 2 and 3) and RNase KO (replicate 2). **C** Box plot representation of data from Figure 3A and S3A and B. **D** Metagene average analysis of trimer 2-5A treated and untreated WT (replicate 2 and 3) and RNase KO (replicate 2) at the main ORF start site. **E-F** Average ribosome footprints around 5’UTR CUG (E) and AUG (F) peaks in trimer 2-5A treated and untreated WT (replicate 2 and 3) and RNase KO (replicate 2) cells. **G** and **H** Increased average ribosome footprints in frame +1 on frame +1 iORFs (G) or −1 (H) on −1 iORFs in 2-5A treated cells in replicate 2 and 3, similar to that of Figure 4F and H. **I** Average ribosome footprints around AUG start codons in the +1 reading frame of coding sequences show a peak in 2-5A treated WT cells. Data from Figure 4F (right panel), reads in the +1 frame only, are plotted (red). As a control here, reads in the −1 frame are also plotted (black) and do not show strong enrichment on +1 AUGs, as expected.

**S4.**
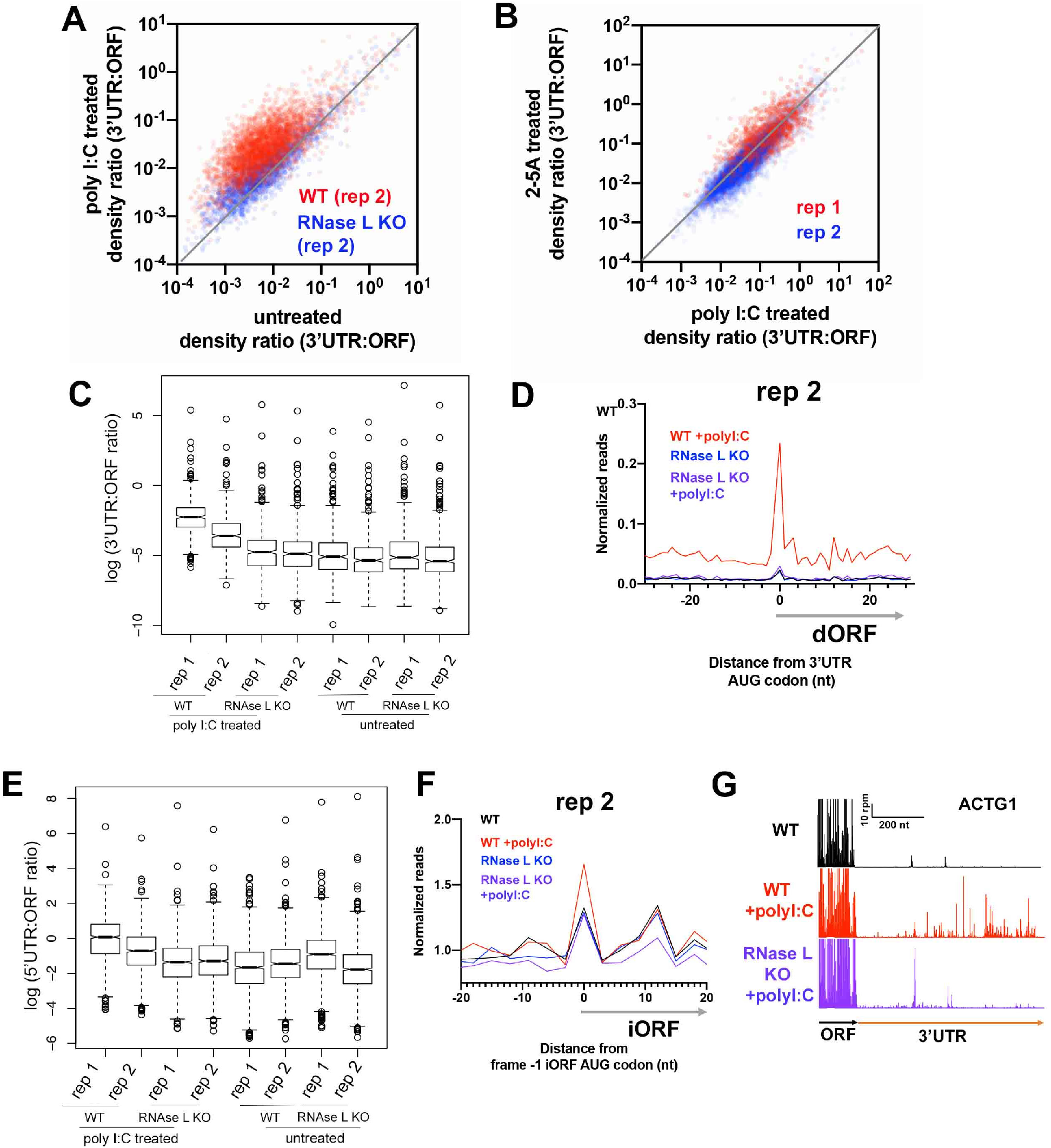
**Related to Figure 5. A** Comparison of 3’UTR:ORF ribosome footprint density ratios in poly I:C treated and untreated WT and RNase KO (replicate 2). Each dot represents one gene model. Genes plotted above the diagonal have increased relative 5’UTR translation in treated in contrast to the untreated sample. All plots include data from genes that passed threshold in both datasets (at least 2 raw reads in both 3’UTRs and in the main ORFs). As discussed in the text, the modest increase in 3’UTR:ORF density ratios in RNase L KO cells (blue) is weaker than when RNase L is present (red) (see full analysis in S4C). **B** Comparison of 3’UTR:ORF ribosome footprint density ratios for 2-5A vs poly I:C treated WT A549 cells (replicate 1 and 2). All plots include data from genes that passed threshold in both datasets (at least 2 raw reads in both 3’UTRs and in the main ORFs). The data are correlated (Pearson’s R^2^=0.60 and 0.85), showing that both treatments evoke a similar 3’UTR translation phenomenon. **C** Box plot representation of data from Figure 4A and S4A. **D** 3’UTR metagene analysis of ribosome footprints showed increased AUG peaks in poly I:C treated WT cells, but not in RNase L KO cells (replicate 2). **E** Box plot representation of data from Figure 4E and additional replicates. **F** Frame −1 ribosome footprints derived from frame −1 internal ORF metagene average plot (generated similarly to Figure 4D) showed increased peaks at iORF AUG start sites. **G** Example gene model of ACTG1 showing increased 3’UTR footprints in poly I:C treated WT cells relative to untreated cells (red vs black). However, we noted a small increase in footprints that follow a different pattern in poly I:C treated KO cells (compare purple to Figure 2C). As discussed in the main text, this likely results from a RNase L independent effect.

**S5.**
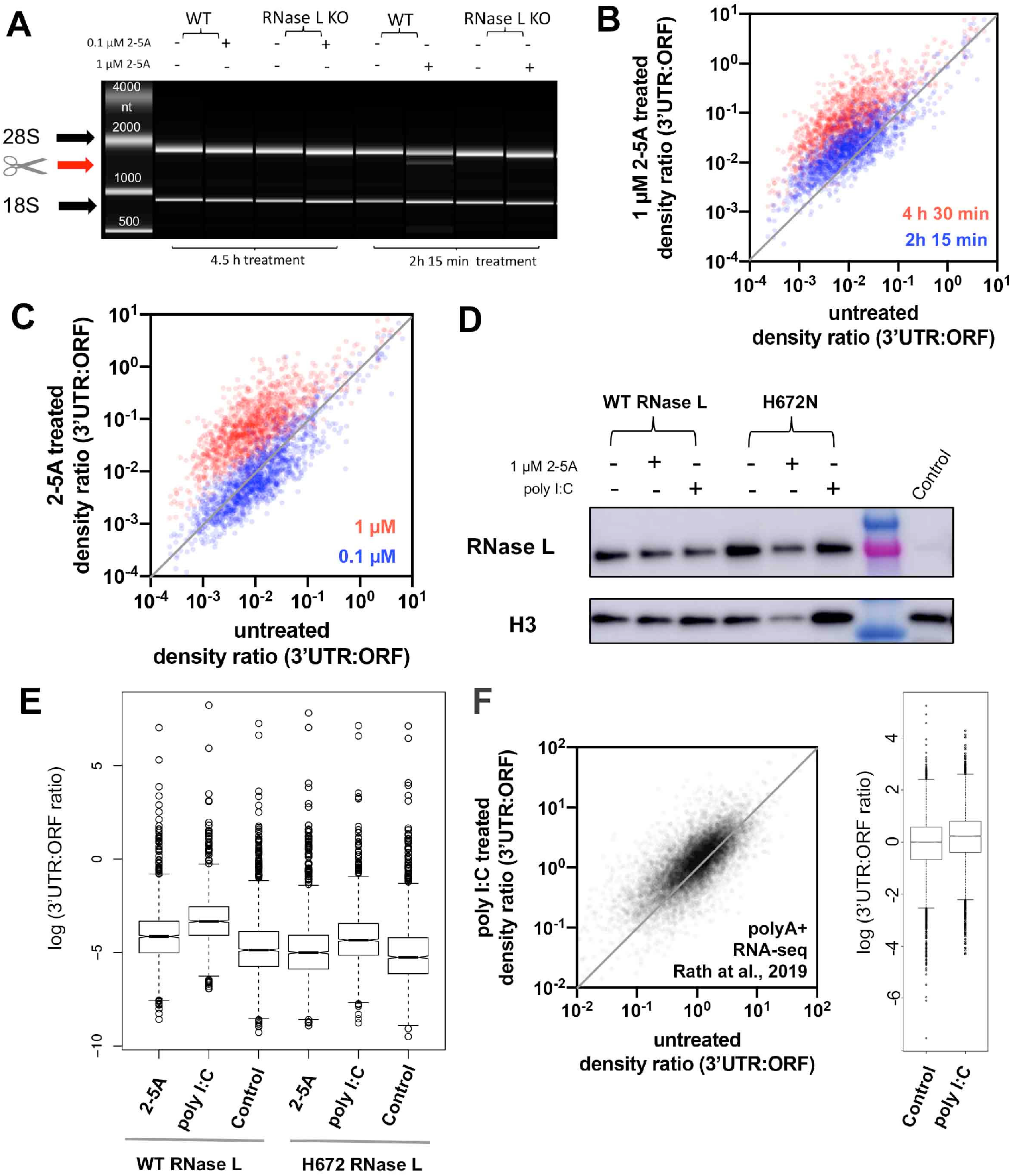
**Related to Figure 6. A** rRNA cleavage assay of 0.1 and 1μM 2-5A treated (treated for 4.5 h and 2h 15 min, respectively) WT and RNase L KO A549 cells. Cleavage products were detected only for 1μM 2-5A treated WT A549 cells (treated for 2h 15 min). **B** Comparison of 3’UTR:ORF density ratios of transcripts in 1 μM 2-5 A treated and untreated (treated for 2h 15 min or 4 h 30 min) WT A549 cells. **C** Comparison of 3’UTR:ORF ribosome footprint density ratios of transcripts in 0.1 or 1 μM 2-5A treated and untreated WT A549 cells. Each dot represents one gene in all dot plots in Figure 6. Genes plotted above the diagonal have increased 3’UTR:ORF ribosome footprint density ratios in treated sample as compared to that of untreated. **D** Expression levels of wild type and H672N RNase L are comparable (within ~2-fold) in all samples used for Ribo-seq. Antibody against Histone H3 was used as a loading control. RNase L KO cells (control) have no detectable RNase L expression. **E** Box plot representation of 3’UTR:ORF ribosome profile density ratios data from Figure 6C and D. **F** Comparison of 3’UTR:ORF polyA^+^ RNA-seq 3’UTR:ORF density ratios for 1 μg/ml poly I:C treated vs untreated A549 cells. Data is processed and analyzed from publicly available polyA^+^ RNA-seq datasets (Accession numbers: SRX5073651 and SRX5073647) (Rath et al., 2019). Genes have increased 3’UTR:ORF ratios in the treated sample as compared to that of untreated, as evidenced by the shift of 3’UTR:ORF density ratios for majority of the genes above the diagonal.

**S6.**
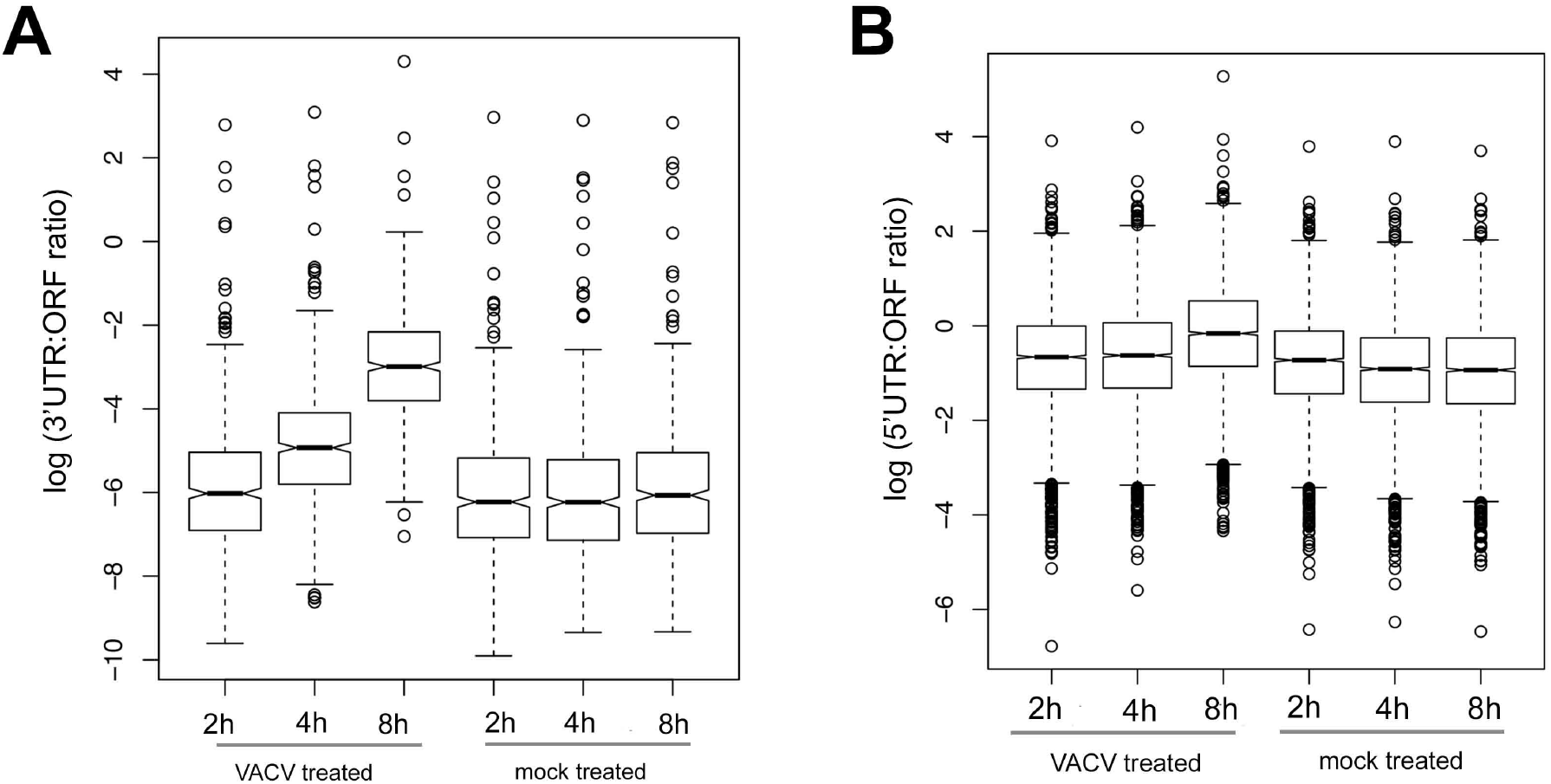
**A Related to Figure 7.** Box plot representation of 3’UTR:ORF density ratios from Figure 7A and other data sets (GEO accession number: SRP056975). **B** Average ribosome footprints around AUG start codons in 5’UTRs show an increased peak at the start codon in 8h VACV treated cells, but not in mock treated or 2 h treated cells. **C** Box plot representation of 5’UTR:ORF ribosome profile density ratios data from Figure 7E and other data sets (GEO accession number: SRP056975).

